# Scaling up spatial transcriptomics for large-sized tissues: uncovering cellular-level tissue architecture beyond conventional platforms with iSCALE

**DOI:** 10.1101/2025.02.25.640190

**Authors:** Amelia Schroeder, Melanie Loth, Chunyu Luo, Sicong Yao, Hanying Yan, Daiwei Zhang, Sarbottam Piya, Edward Plowey, Wenxing Hu, Jean R. Clemenceau, Inyeop Jang, Minji Kim, Isabel Barnfather, Su Jing Chan, Taylor L. Reynolds, Thomas Carlile, Patrick Cullen, Ji-Youn Sung, Hui-Hsin Tsai, Jeong Hwan Park, Tae Hyun Hwang, Baohong Zhang, Mingyao Li

**Affiliations:** Statistical Center for Single-Cell and Spatial Genomics, Department of Biostatistics, Epidemiology and Informatics, Perelman School of Medicine, University of Pennsylvania, Philadelphia, PA 19104, United States; Departments of Biostatistics and Genetics, University of North Carolina, Chapel Hill, NC 27599, United States; Research Department, Biogen Inc., Cambridge, MA 02142, United States; Department of Surgery, Vanderbilt University Medical Center, Nashville, TN 37232, United States; Department of Pathology, College of Medicine, Kyung Hee University Hospital, Kyung Hee University, Seoul, Republic of Korea; Department of Pathology, Seoul Metropolitan Government-Seoul National University Boramae Medical Center, Seoul National University College of Medicine, Seoul, Korea; Department of Pathology and Laboratory Medicine, Perelman School of Medicine, University of Pennsylvania, PA 19104, United States

**Keywords:** spatial transcriptomics, super-resolution, histology, machine learning

## Abstract

Recent advances in spatial transcriptomics (ST) technologies have transformed our ability to profile gene expression while retaining the crucial spatial context within tissues. However, existing ST platforms suffer from high costs, long turnaround times, low resolution, limited gene coverage, and small tissue capture areas, which hinder their broad applications. Here we present iSCALE, a method that predicts super-resolution gene expression and automatically annotates cellular-level tissue architecture for large-sized tissues that exceed the capture areas of standard ST platforms. The accuracy of iSCALE were validated by comprehensive evaluations, involving benchmarking experiments, immunohistochemistry staining, and manual annotation by pathologists. When applied to multiple sclerosis human brain samples, iSCALE uncovered lesion associated cellular characteristics that were undetectable by conventional ST experiments. Our results demonstrate iSCALE’s utility in analyzing large-sized tissues with automatic and unbiased tissue annotation, inferring cell type composition, and pinpointing regions of interest for features not discernible through human visual assessment.

## Introduction

Understanding spatial distributions for gene expression profiles across tissues is essential for answering crucial biological questions and advancing our knowledge of cellular functions, disease mechanisms, and tissue development. Spatial transcriptomics (ST) technologies have transformed our ability to profile gene expression across tissues while retaining crucial spatial context^1–3^. However, a significant barrier to the widespread adoption of commercial ST platforms in biomedical research is their high costs, long turnaround times, low resolution, limited gene coverage, and small tissue capture area offered by most ST platforms. Sequencing based ST platforms, such as Visium^4^, can sequence the whole transcriptome, but lack single-cell resolution and come with a standard tissue capture area of 6.5 mm × 6.5 mm. While Visium offers an extended version with a larger capture area of 11 mm × 11 mm, it is more expensive than the standard version. Furthermore, many tissue samples still surpass this size limitation, making it unable to capture entire biological entities in a single capture. The recently released Visium HD ^5^ offers subcellular resolution, but its cost is significantly higher than Visium, and its tissue capture area is only 6.5 mm × 6.5 mm. Imaging-based ST platforms like MERSCOPE^6^, CosMx^7^, and Xenium^8^ provide subcellular resolution and can handle larger-sized tissues, but the number of genes is limited and image scanning time is typically extensive. For instance, in the current version of the CosMx platform, scanning a 200 mm^2^ tissue area with ~1000 genes takes about seven days and the scanning time increases linearly as the tissue size and number of field of views increase. These limitations make commercial ST platforms impractical for large-scale investigations, particularly when studying sizable tissues, which is common in human studies.

In contrast, hematoxylin and eosin (H&E)-stained histology images, routinely used by clinical pathology labs, are considerably more cost-effective than ST. Additionally, the physical size of a typical whole slide H&E image can be as large as 25 mm × 75 mm, greatly exceeding the capture area of all ST platforms. Previous studies have shown correlations between gene expression profiles and histological image characteristics^9–13^, suggesting the feasibility of inferring spatial gene expression from these histology images^14–18^. Since obtaining a histology image from a large-sized tissue is easily achievable, this opens the possibility of virtually predicting gene expression in such tissues directly from histology images. This approach makes it possible to generate virtual ST data, overcoming the tissue capture area size limitations of conventional ST platforms.

In this paper, we introduce iSCALE (**i**nferring **S**patially resolved **C**ellular **A**rchitectures in **L**arge-sized tissue **E**nvironments), a novel machine learning framework designed to predict gene expression for large-sized tissues with cellular resolution. By leveraging the gene expression-histological feature relationship, learned from a small set of training ST captures, iSCALE enables comprehensive gene expression prediction and cellular-level tissue annotation across entire large tissue sections, including tissue regions where gene expression has not been directly measured. Our results demonstrate that iSCALE can provide critical insights into disease by generating high-resolution spatial gene expression maps for large tissue sections. These maps encompass entire diseased biological entities of interest, along with surrounding immune and adjacent normal tissue regions. iSCALE has the potential to advance biomedical research and contribute to progress in the understanding, diagnosis, and treatment of human disease.

## Results

### Overview of iSCALE

An overview of the iSCALE workflow is shown in **Fig. 1**. Our goal is to predict gene expression in an H&E image generated from a large-sized tissue section, referred to as the ‘mother image’. The pipeline begins by selecting regions from the same tissue block, potentially across multiple tissue sections, with region sizes fitting the capture areas of standard ST platforms (**Supplementary Fig. 1**). This process generates a small set of training ST data, referred to as ‘daughter captures’. iSCALE then implements clustering analysis on the daughter ST data to guide the alignment of the daughter captures onto the mother image through a human-in-the-loop, semi-automatic process (**Supplementary Fig. 2**). Next, iSCALE harmoniously integrates the gene expression and spatial information across the aligned daughter captures. To construct a gene expression prediction model, iSCALE extracts both global and local tissue structure information from the mother image. Subsequently, a feed-forward neural network is employed to learn the relationship between histological image features and gene expression transferred from the aligned daughter captures. The resulting predictive model predicts gene expression for each 8 µm × 8 µm superpixel—corresponding to approximately the size of a single cell—across the entire mother image. Leveraging these predictions, iSCALE further annotates each superpixel in the mother image with cell types, identifies enriched cell types in each tissue region, providing a detailed characterization of tissue architecture at cellular resolution.

**Fig. 1.**
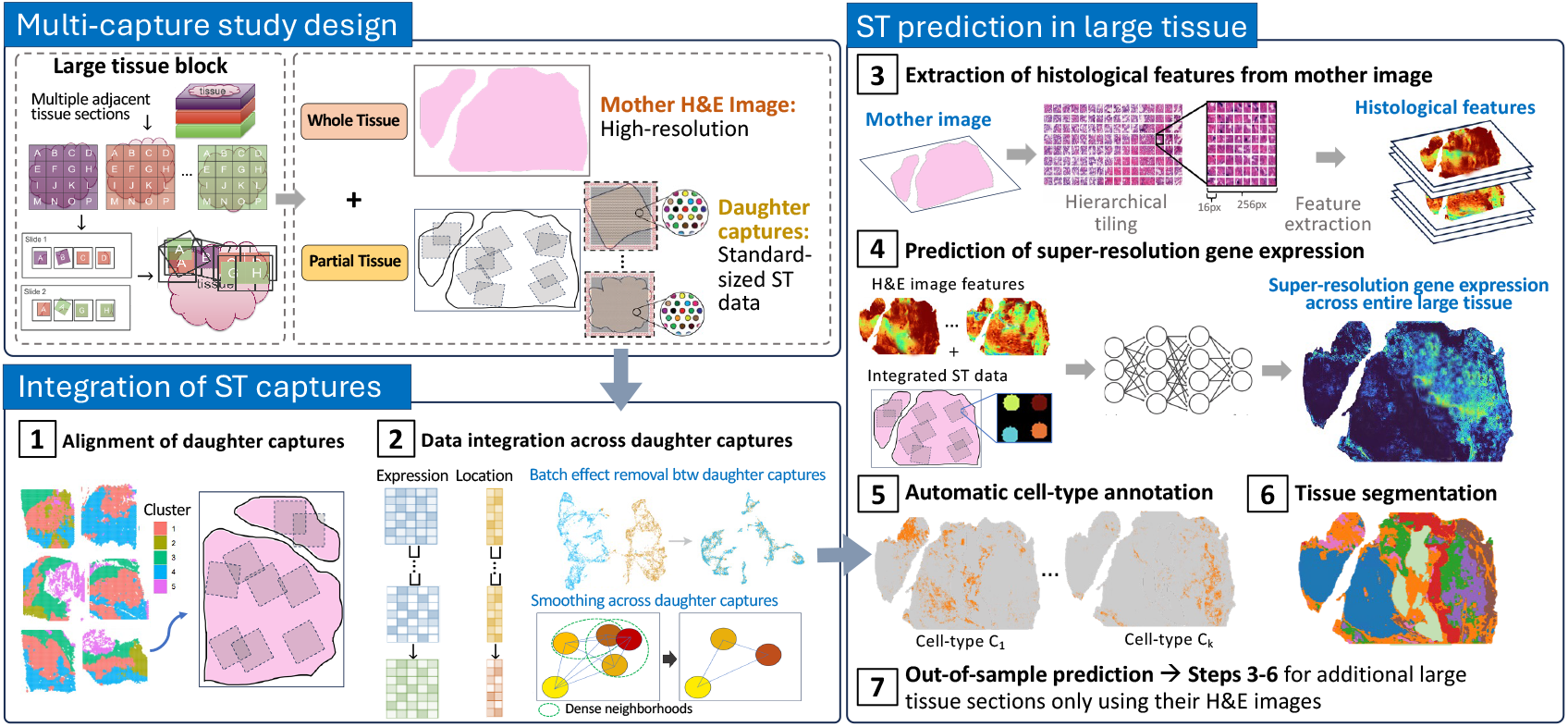
Overview of iSCALE workflow. iSCALE operates on a study design where adjacent tissue slices are obtained from a large tissue block. A high-resolution mother H&E image is generated from one slice, while multiple smaller daughter captures, containing both gene expression and spatial location information, are obtained from adjacent slices. These smaller daughter captures are clustered based on their gene expression and aligned to the large mother H&E image. The mother image is hierarchically divided into tiles, which are then converted into histology image features. These features, combined with integrated spot-level gene expression data from the daughter captures, are used to predict super-resolution gene expression across the entire large tissue section, including regions not originally covered by the daughter captures. Using the predicted super-resolution gene expression, iSCALE infers detailed cell types and tissue architecture for the entire section. Additionally, out-of-sample predictions can be performed on other large H&E tissue sections.

### Benchmark evaluation of iSCALE on a large gastric cancer sample

To evaluate the effectiveness of iSCALE, we conducted a benchmarking experiment using a ground truth single-cell gene expression dataset derived from a large tissue sample. This experiment was obtained from a gastric cancer tissue section measured with the 10x Xenium platform, encompassing 377 genes. The tissue section spans the entire Xenium slide (12 mm × 24 mm), making it an ideal choice for benchmarking. In this experiment, we simulated a scenario where gene expression data were available only from five daughter captures, each with size of 3.2 mm × 3.2 mm, mimicking the conditions typically observed in real studies (**Fig. 2a**). Within each daughter capture, we simulated pseudo-Visium data, following the spot size and layout of the Visium platform. The iSCALE model was then trained using the integrated gene expression data from the five daughter captures, assuming their true alignment on the mother image. For comparison, we also predicted gene expression using iStar^14^ and RedeHist^17^. Since both iStar and RedeHist can process only one daughter capture at a time to build their prediction models, we applied these algorithms for each daughter capture.

**Fig. 2.**
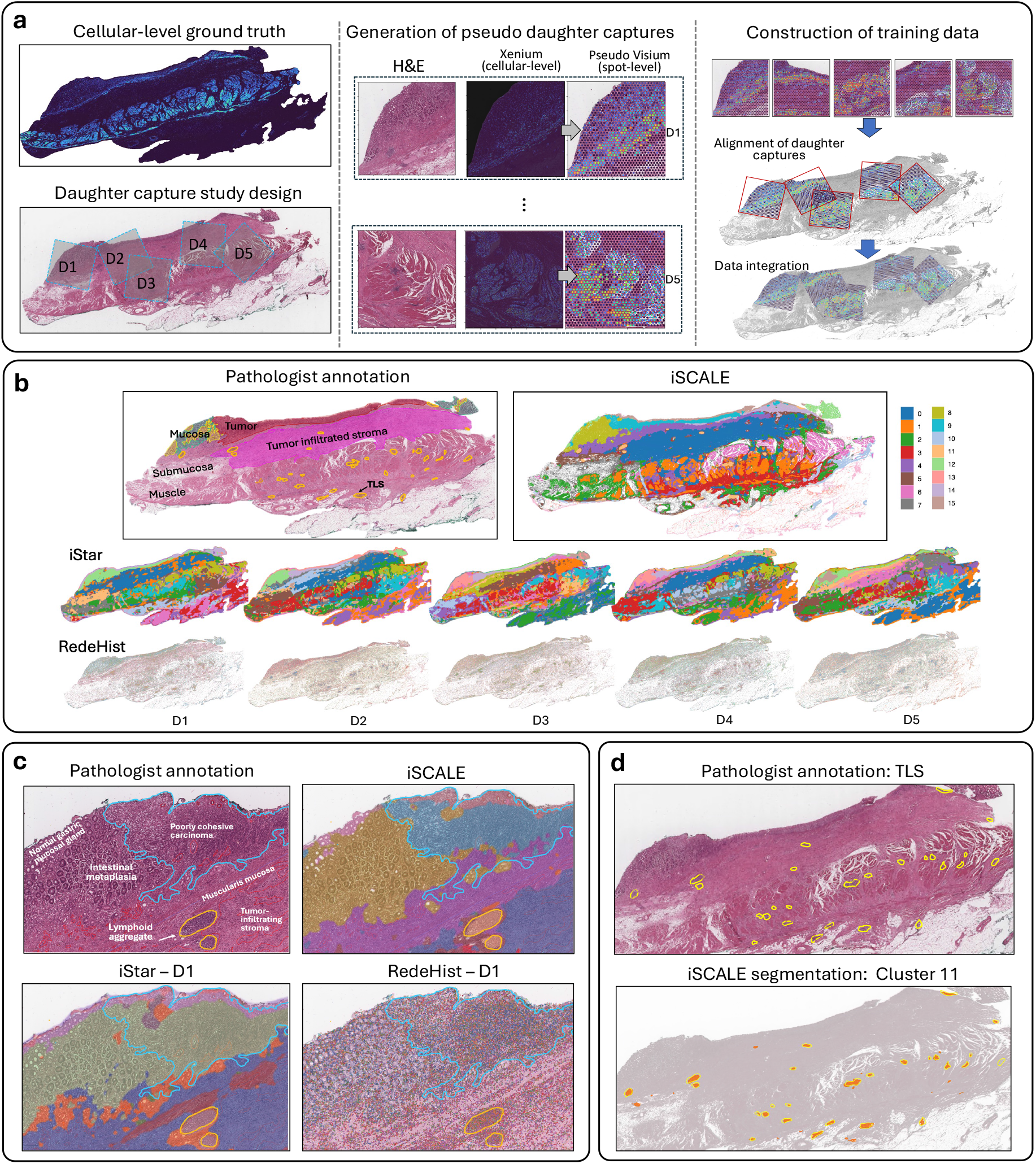
Evaluation of iSCALE segmentation in the gastric cancer benchmarking data. **a**, Overview of the benchmarking design. Ground truth gene expression data were obtained using 10x Xenium. Five daughter captures, each measuring 3.2 mm × 3.2 mm, were generated with pseudo-Visium gene expression data. These daughter captures were aligned and integrated to generate a training dataset for the iSCALE model. **b**, Pathologist manual annotation, and tissue segmentation using the predicted gene expressions from iSCALE, iStar, and RedeHist. **c**, Detailed examination of the upper-left region of the tissue. For iStar and RedeHist, the segmentation results from the D1-trained model are displayed, as D1 is the only daughter capture among the five that covers the boundary between the tumor and mucosa. **d**, TLSs annotated by pathologist and detected by iSCALE.

**Fig. 2b** shows the tissue segmentation results obtained from iSCALE, iStar, and RedeHist. iSCALE’s segmentation closely resembles the pathologist’s manual annotation, and successfully identified key tissue structures, including tumor, tumor infiltrated stroma, mucosa, submucosa, muscle, and tertiary lymphoid structure (TLS). In contrast, noticeable differences were observed among the five segmentations made by both iStar and RedeHist, depending on which daughter capture was used for model training. For instance, iStar struggled to distinguish between tumor and mucosa when using daughter captures D1 and D5 as the training data. RedeHist performed even more poorly, failing to identify tissue structures regardless of which daughter capture was used as the training data. This is likely due to its inability to correctly detect a large portion of the nuclei (**Supplementary Fig. 3**), a critical component of its algorithm. To further demonstrate the advantage of iSCALE, we focused on the upper-left region of the tissue containing signet ring cells (**Fig. 2c**), which are associated with aggressive gastric cancer, poor prognosis, and play a critical role in guiding treatment^19^. iSCALE accurately identified the boundary between the poorly cohesive carcinoma region with signet ring cells and the adjacent gastric mucosa, with its inferred boundary closely aligning with the pathologist’s manual annotation. However, both iStar and RedeHist failed to detect this boundary when using daughter capture D1, which covers that boundary for model training (**Fig. 2c; Supplementary Fig. 4**). As another demonstration, we examined the TLSs in this sample. Detecting TLSs in cancer is crucial because they are associated with improved immune responses, enhanced anti-tumor activity, and better patient prognosis, making them key indicators of the tumor microenvironment’s immune dynamics^20–22^. As shown in **Fig. 2d**, cluster 11 (peach color) identified by iSCALE closely aligns with the manually annotated TLSs. In contrast, iStar tends to detect false positive TLSs, whereas RedeHist exhibited significantly lower detection accuracy (**Fig. 2d; Supplementary Fig. 5**). Notably, daughter captures that performed well for certain annotations often performed poorly for others critical regions, demonstrating significant variability among daughter capture predictions. These results underscore the importance of integrating multiple daughter captures to accurately detect fine-grained tissue structures such as TLSs.

The availability of the ground truth single-cell gene expression data in this benchmarking dataset enables a quantitative evaluation of iSCALE’s gene expression prediction accuracy. Although iSCALE was designed for Visium data, its prediction model can be readily adapted to use Xenium data for training. In the following evaluations, we compared two versions of iSCALE: iSCALE-Seq, which used pseudo-Visium data for training, and iSCALE-Img, which used Xenium data for training. To evaluate the prediction performance, we focused on the top 100 highly variable genes. We computed the following metrics, including RMSE (root mean squared error), SSIM (structural similarity index measure), and Pearson Correlation. As shown in **Fig. 3a,b**, iSCALE-Seq outperformed iStar across all evaluation metrics, achieving performance comparable to iSCALE-Img, despite being trained on spot-level resolution ST data. RedeHist was excluded from this comparison due to its unsatisfactory performance (**Supplementary Fig. 6**). Although the Pearson Correlation coefficients for iSCALE-Img and iSCALE-Seq were generally low at the superpixel level (8 µm × 8 µm), the correlations improved as the superpixel size increased. Moreover, we observed low correlation values, such as r=0.27 for *ACTA2*, still resulted in spatial distributions that almost perfectly resembled the ground truth predictions (**Fig. 3c**). Approximately 50% of the genes achieved correlation coefficients above 0.45 at a spatial resolution of 32 µm × 32 µm. For illustration purpose, we visualized the predicted gene expression levels obtained from iSCALE and iStar, comparing with the ground truth single-cell gene expression directly measured using Xenium (**Fig. 3c; Supplementary Fig. 6**). Across all examined genes, iSCALE’s predictions showed the closest resemblance to the ground truth gene expression. For instance, for gene *TFF2*, iStar’s predictions exhibited a strong dependence on the specific daughter capture used for training. Additionally, the optimal daughter capture for iStar predictions varied across different genes. This highlights the importance of integrating data from all daughter captures when constructing the prediction model, a strength clearly demonstrated by iSCALE.

**Fig. 3.**
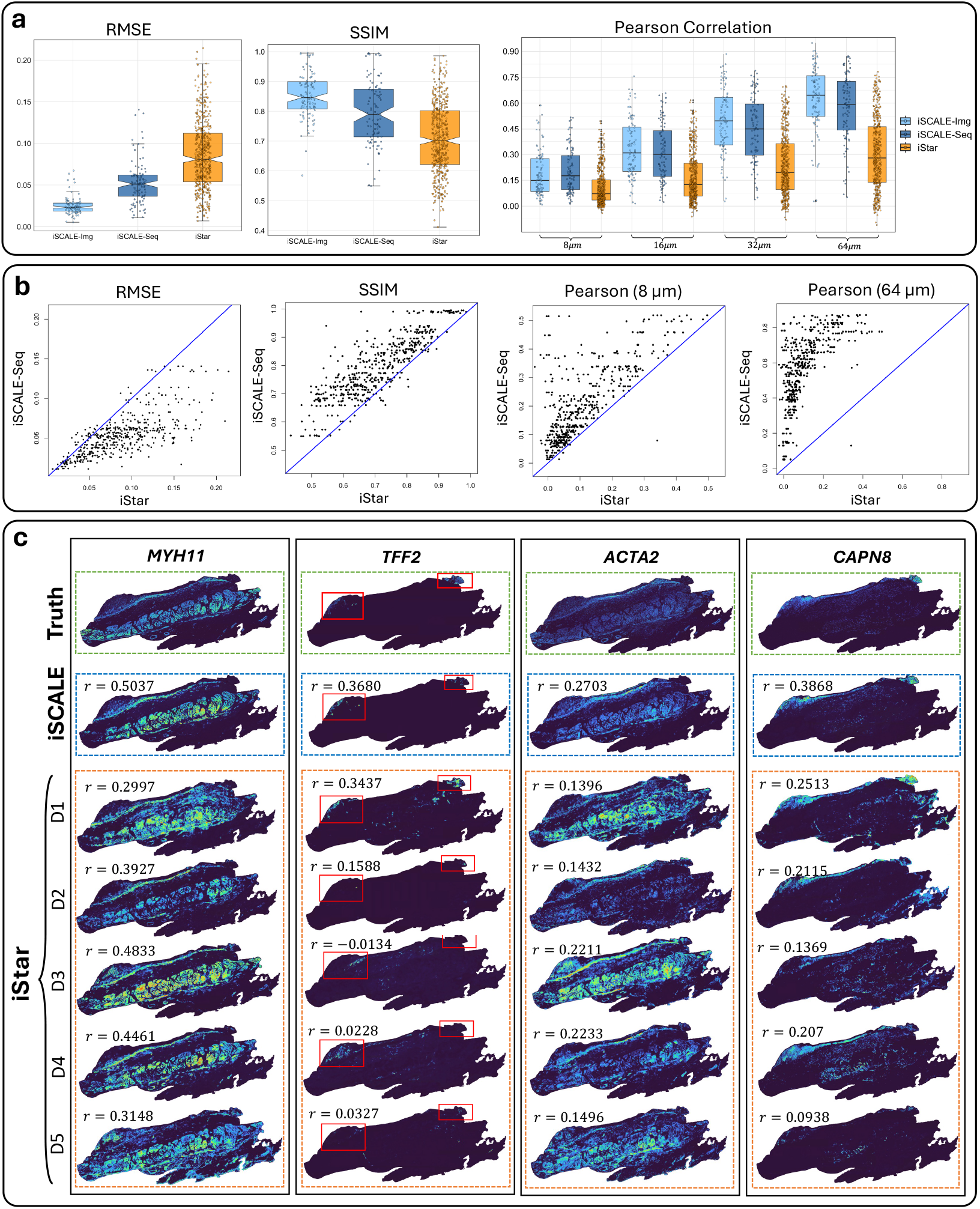
Evaluation of iSCALE prediction accuracy in the gastric cancer benchmarking data. **a**, Quantitative comparison of iSCALE and iStar predictions by RMSE, SSIM, and Pearson Correlation based on the top 100 highly variable genes. RMSE and SSIM calculations were calculated at a spatial resolution of 8 µm, while Pearson Correlation was evaluated across multiple spatial resolutions. iSCALE-Seq: iSCALE trained using the pseudo-Visium data in the daughter captures. iSCALE-Img: iSCALE trained using the Xenium data in the daughter captures. The hinges are the boundaries of the box corresponding to the 25th percentile and 75^th^ percentile of the data. The whiskers extend to data points within 1.5 × interquartile range from the hinge. Data beyond the end of the whiskers are plotted individually. **b**, Comparison of iSCALE-Seq and iStar predictions at the per-gene level for the top 100 highly variable genes. **c**, Visualization of ground truth gene expression directly measured by Xenium, and predicted super-resolution gene expression by iSCALE and iStar at the 8 µm spatial resolution. The red box highlights where *TFF2* was expressed in the ground truth data.

In the above evaluations, we assumed the true alignment of the daughter captures on the mother image. To assess the accuracy of iSCALE’s semi-automatic alignment algorithm, we aligned the five daughter captures using the proposed algorithm described in Methods. As shown in **Supplementary Fig. 7**, the alignment achieved an accuracy of 99%. This high accuracy is likely due to the relative simplicity of the benchmarking scenario compared to real-world data. As demonstrated in later sections, we applied this alignment algorithm to more challenging human brain data, where the daughter captures and the mother image were derived from different tissue sections. Despite differences in tissue shape, iSCALE successfully aligned the daughter captures to their expected positions on the mother image.

### Benchmark evaluation of iSCALE on two normal gastric samples

Next, we evaluated iSCALE’s performance for out-of-patient prediction, where the test sample was from a different patient and included only an H&E image with no available gene expression data. For this benchmarking experiment, we used two normal gastric tissue sections obtained from different patients, both measured using the 10x Xenium platform with 377 genes. Each tissue section was relatively large, spanning the entire Xenium slide. To simulate the scenario (**Fig. 4a**), we assumed that gene expression data were available only from 10 daughter captures (each 2 mm × 2 mm) in the training sample (Normal 1), while the test sample (Normal 2) had no gene expression data. We also simulated eight daughter captures (each 2 mm × 2 mm) in Normal 2 to compare iSCALE’s out-of-patient prediction with its in-sample prediction for Normal 2. As shown in **Fig. 4b**, the segmentation result from the out-of-patient prediction closely aligns with that from the in-sample prediction and pathologist annotation, achieving an Adjusted Rand Index (ARI) of 0.74 when comparing the two segmentations.

**Fig. 4.**
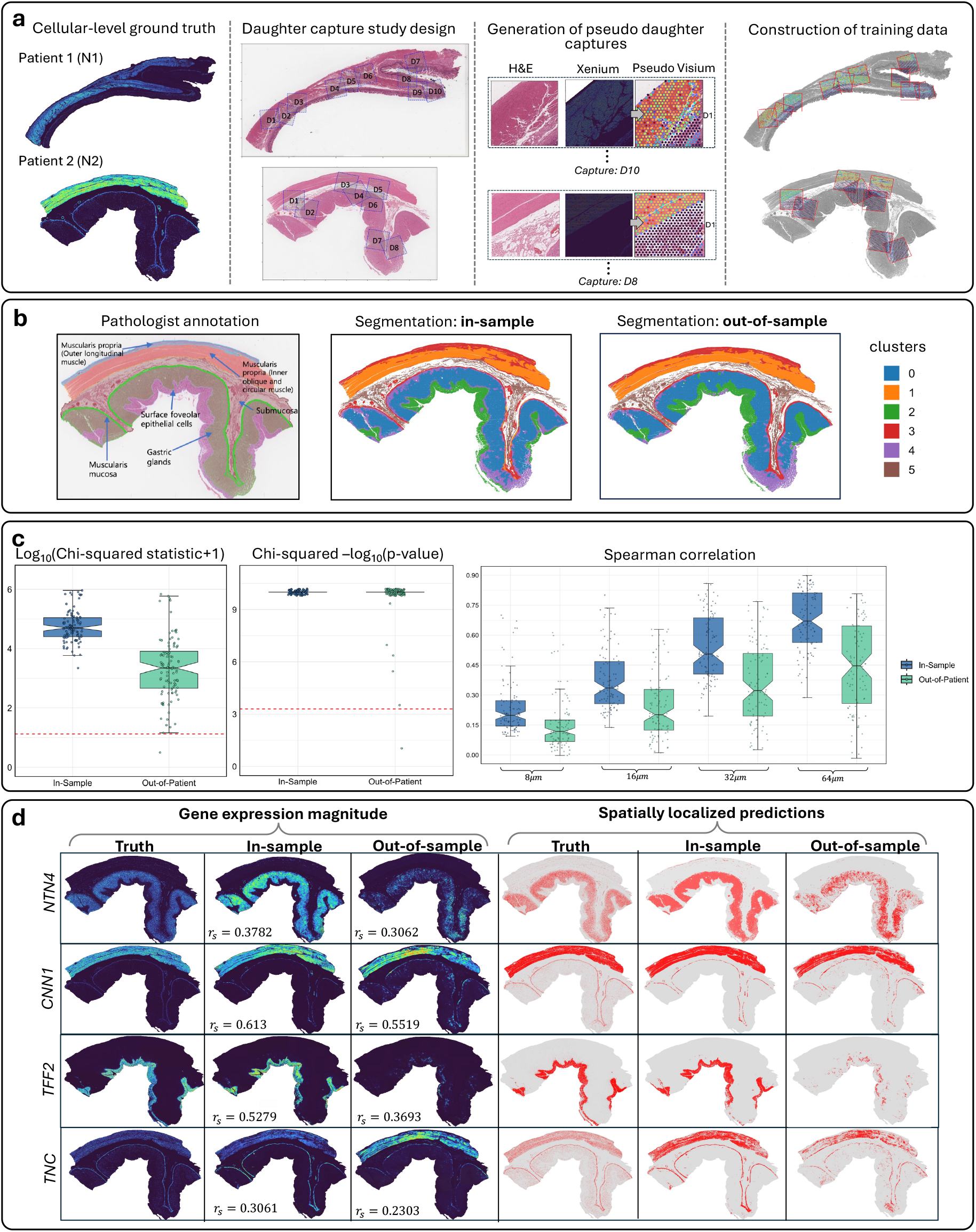
Evaluation of iSCALE segmentation and prediction accuracy in the normal gastric tissue benchmarking data. **a**, Overview of the benchmarking design. Ground truth gene expression data were obtained using 10x Xenium. Ten daughter captures for Normal 1 and eight daughter captures for Normal 2, each measuring 2 mm × 2 mm, were generated with pseudo-Visium gene expression data. These daughter captures were aligned and integrated to generate a training dataset for the iSCALE model. **b**, Pathologist manual annotation, and tissue segmentations using the predicted gene expressions from the in-sample and out-of-sample analyses. For in-sample predictions, the iSCALE model was trained using the eight daughter captures from Normal 2. For out-of-sample predictions, the iSCALE model was trained using the 10 daughter captures from Normal 1. **c**, Quantitative comparison of iSCALE in-sample and out-of-sample predictions by Chi-squared statistic and Spearman Correlation for the top 100 highly variable genes. Chi-squared statistic was calculated at a spatial resolution of 8 µm, while Pearson Correlation was evaluated across multiple spatial resolutions. For the Chi-squared statistic, the red dashed lines indicate significance thresholds set at the 0.05/100 level. The hinges are the boundaries of the box corresponding to the 25th percentile and 75^th^ percentile of the data. The whiskers extend to data points within 1.5 × interquartile range from the hinge. Data beyond the end of the whiskers are plotted individually. **d**, Visualization of ground truth gene expression directly measured by Xenium, and predicted super-resolution gene expression by iSCALE in-sample and out-of-sample predictions, shown at the 8 µm spatial resolution. The visualizations include the predicted gene expression magnitude and the corresponding binary predictions.

We further evaluated the accuracy of gene expression predictions for the top 100 highly variable genes. Since the range of gene expression is unobserved for out-of-patient predictions, evaluation metrics such as RMSE, SSIM, and Pearson Correlation, which rely on the magnitude of gene expression, are not suitable. Instead, we utilized Spearman Correlation, which is rank-based, and introduced a Chi-squared statistic to assess concordance between binarized gene expression patterns from the ground truth and predictions. Gene expression was binarized as either ‘expressed’ or ‘not expressed’, allowing for a direct comparison of overall expression patterns using the Chi-squared statistic. As shown in **Fig. 4c**, of the 100 genes, 99 exhibited significantly concordant predicted expression patterns with the ground truth. While the correlations for out-of-sample predictions were lower than those for in-sample predictions, at the spatial resolution of 64 µm, approximately 50% of the genes achieved correlation coefficients greater than 0.45. **Fig. 4d** and **Supplementary Fig. 8** show a few representative genes where iSCALE’s out-of-patient gene expression predictions resemble those from the in-sample predictions. While predicting gene expression magnitude is challenging, the out-of-patient predictions achieved reasonable correlation coefficients when compared with in-sample predictions. Furthermore, when focusing on whether a superpixel is expressed or not, the out-of-patient predictions closely matched in-sample predictions for three out of the four genes examined.

### Application of iSCALE to a large postmortem human brain sample with multiple sclerosis

Encouraged by the promising performance of iSCALE in benchmark evaluations, we next applied it to data generated from a postmortem human brain of a multiple sclerosis (MS) patient. MS is a chronic inflammatory disorder that affects the central nervous system (CNS) by mistakenly attacking the myelin sheath of nerve fibers, leading to inflammation and damage, which disrupts electrical impulse transmission along nerves^23–25^. Our focus is on cellular alterations in MS, specifically changes in abundance, distribution, or activity of various cell types within and near MS lesions, including CNS cell types such as microglia, astrocytes, oligodendrocytes, and neurons. Identifying distinct cellular profiles around MS lesions can offer insights into the disease’s pathology, the immune response it provokes, and potential therapeutic targets.

To characterize cell attributes associated with MS lesions, we analyzed a tissue block (Sample 1) containing a complete white matter inactive lesion core from an MS patient. Since the size of the mother tissue section (~22 mm × 19 mm = 418 mm^2^) exceeded the standard ST capture area (**Supplementary Fig. 9a**), we obtained 11 smaller daughter captures from the same tissue block following the study design in **Fig. 1**. These captures underwent gene expression analysis using the standard Visium protocol with tissue capture area of 6.5 mm × 6.5 mm (**Supplementary Table 1**). Using the alignment process described in the Methods section, we aligned these daughter captures onto the mother image (**Fig. 5a; Supplementary Fig. 2**). Traditional ST analysis, which relies solely on observed spot-level gene expression from directly measured daughter captures would be inadequate due to its low-resolution, spot-level segmentation, and limited coverage of critical regions such as the lesion rim and core. In contrast, iSCALE integrates gene expression data from all 11 daughter captures and their corresponding regions in the mother image to train a gene expression prediction model for super-resolution gene expression prediction across the entire tissue section. For this analysis, iSCALE predicted the expression of the top 3,000 highly variable genes, along with 145 additional key cell-type markers selected based on in-house single-nucleus RNA-seq data generated from the same brain. As shown in **Supplementary Fig. 10**, the iSCALE pipeline successfully removed gene expression batch effects across all daughter captures. In total, iSCALE predicted super-resolution gene expression across the entire mother image, consisting of 2,013,871 superpixels, each with size of 8 µm × 8 µm. These super-resolution gene expression data allowed iSCALE to segment the tissue and identify regions closely matching pathologist’s manual annotation (**Fig. 5a**). iSCALE successfully detected the white matter inactive lesion core (purple cluster), the rim with increased microglial activation (red cluster), normal appearing white matter (orange cluster), and normal appearing grey matter (blue and green clusters) continuously across the whole mother image.

**Fig. 5.**
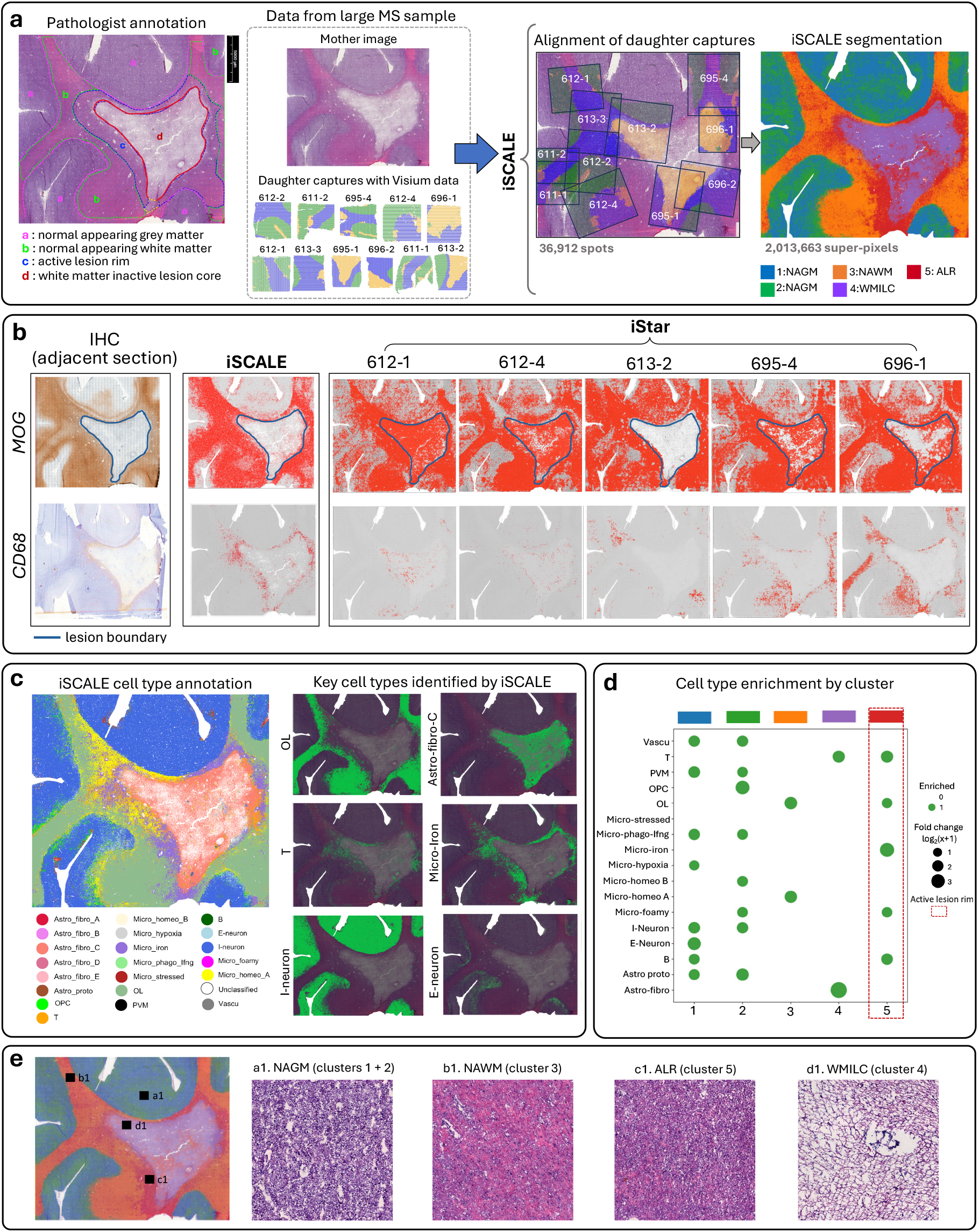
Application of iSCALE to a large postmortem human brain sample with MS. **a**, Mother H&E image with pathologist manual annotation, observed Visium data from 11 daughter captures obtained from adjacent tissue sections, alignment of daughter captures onto the mother image, and segmentation using iSCALE’s predicted gene expressions. Abbreviations: NAGM, normal appearing grey matter; NAWM, normal appearing white matter; WMILC, white matter inactive lesion core; ALR: active lesion rim. **b**, Experimental validation of iSCALE’s predicted gene expression by IHC staining, where the IHC stains were taken from adjacent tissue sections for critical markers including MOG and CD68. For iSCALE and iStar, the binarized gene expression is shown, where a superpixel was considered expressed if its expression exceeded 25^th^ percentile of all MOG predictions and 25the percentile of all CD68 positive predictions. Different thresholds were used because MOG’s expression range is much larger than that of CD68. Since the expression values were standardized to the range [0, 1], using a lower threshold for MOG ensures that the absolute thresholds are more comparable between MOG and CD68. **c**, Cell type annotation of Sample 1 using the predicted super-resolution gene expression from iSCALE with all cell types across the whole mother capture. Abbreviations: OL, oligodendrocyte; I-neuron: inhibitory neuron; E-neuron: excitatory neuron; Astro-fibro-C: fibrous astrocyte subpopulation C; Micro-Iron, iron rich microglia. **d**, Cell type enrichment analysis for the identified clusters from iSCALE. Outlined in red is the enrichment for cluster identified as the hypercellular rim containing reactive microglia by pathologist (E.P.). **e**, Display of heterogeneity in histology across different anatomical regions of the tissue. Shown is an overlay of the iSCALE segmentation result on top of the mother H&E image. The right panel includes a close-up visualization of the high-resolution H&E image for four specific areas within the given biological regions of the tissue corresponding to clusters identified by iSCALE.

Next, we assessed iSCALE’s prediction accuracy using Immunohistochemistry staining (IHC), an important auxiliary method for pathologists. IHC allows the visualization of the distribution and quantity of specific molecules within tissue^16^. This technique was used to validate iSCALE’s predictions for critical markers essential for the identification and classification of MS lesions. Specifically, we investigated the following two markers: MOG (used for discerning MS lesion types based on the relative absence of MOG immunoreactivity, indicating demyelinating or demyelinated regions within the tissue), and CD68 (an activation marker of monocyte lineage). The iSCALE predictions agreed with the IHC staining patterns obtained from adjacent tissue sections as seen in **Fig. 5b**. Consistent with the IHC staining and the demyelinated nature of the MS lesion, the super-resolution gene expression predicted by iSCALE shows negligible expression of MOG within the chronic white matter lesion. Furthermore, iSCALE’s predictions indicate the presence of CD68 expression along the annotated rim of the lesion border, a pattern undetected by iStar using any of the representative daughter captures. Overall, iStar failed to generate accurate predictions for both markers with large variability across captures. For an in-depth analysis, we compared the observed spot-level gene expression and the predicted super-resolution gene expression for cell-type and MS related marker genes. We illustrate this agreement for the following genes: *PLP1* (white matter, oligodendrocyte marker), *SLC17A7* (gray matter, neuron marker), and *HLA-DRA* (active immune response, microglia marker), along with additional MS-related and cell-type marker genes (**Supplementary Fig. 11**). The predicted expression aligned closely with the spatial distribution observed in the tissues while providing significantly enhanced spatial resolution and comprehensive tissue coverage.

We further show that the super-resolution gene expression predictions by iSCALE can provide deeper insights into the cellular attributes associated with MS lesions, thereby facilitating additional downstream cellular level analyses. Using the super-resolution gene expression prediction as input, iSCALE automatically annotated cell types for each superpixel (**Fig. 5c; Supplementary Fig. 12**). We observed distinct spatial distributions of different cell types across the normal and diseased portions of the tissue, including a high density of oligodendrocytes (OL) in the normal-appearing white matter, inhibitory neurons (I-neuron) and excitatory neurons (E-neuron) in grey matter, fibrous astrocytes subpopulation C (Astro-fibro C) within the inactive lesion core, and a mixture of iron rich microglial (Micro-iron) and T cells in the lesion rim (**Fig. 5c**). By inferring cell types within these superpixels, we gained insights into the spatial distribution and diversity of cell populations within the tissue, allowing for comprehensive analysis and interpretation of the biological landscape. This is particularly valuable in regions of tissue that are challenging to differentiate through histological inspection or gene expression profiling alone. For example, the red cluster, corresponding to the active lesion rim, is enriched with immune response-related cell types, including T cells, B cells, and several microglia subpopulations such as iron rich microglia and foamy microglia (micro_foamy), along with infiltrated T cells within the lesion. These subpopulations are challenging for pathologists to discern from H&E images. As a comparison, we performed cell type annotation using iStar-predicted gene expressions. Unsurprisingly, the annotations were highly variable depending on which daughter capture was used for model training, and the cells were frequently annotated in locations that did not align with biological expectations. For instance, when daughter capture 695-4 was used for model training, iStar predicted inhibitory neurons in the inactive lesion core, a result that contradicts established knowledge of MS lesion biology (**Supplementary Fig.13**).

To further validate the biological relevance of the iSCALE segmentation results, we focused on key comparisons among the following main tissue regions annotated by the pathologist, including normal appearing grey matter (clusters 1 and 2, region a), normal appearing white matter (cluster 3, region b), active lesion rim (cluster 5, region c), and white matter inactive lesion core (cluster 4, region d) (**Fig 5e**). Between these distinct regions, we examined the corresponding H&E image by selecting representative image patches from each of the clusters. A noticeable difference in tissue structure can be observed in the H&E image between these three regions as shown in **Fig 5e**. In contrast to the highly fibrous clusters found within the inactive lesion core, the active lesion rim presents a denser, heterogeneous structure characterized by a mixture of various immune cells. Upon close examination of the H&E images corresponding to the iSCALE cluster comprising the active lesion rim (cluster 5), pathologist (E.P.) confirmed that this cluster clearly corresponds to the hypercellular rim containing reactive microglia. Moreover, the pathologist (E.P.) observed that the normal appearing white matter region (cluster 3, region b) discovered by iSCALE, corresponds to less smoldering microglial neuroinflammation compared to the active lesion rim.

### Application of iSCALE to an additional large postmortem human brain MS sample

In this section, we explore the feasibility of using the prediction model trained on Sample 1 to predict gene expression in another MS sample. The test sample, Sample 2, was obtained from a different region of the same MS brain as Sample 1. Sample 2 contains a complete white matter chronic active lesion and areas of active and chronic subpial cortical demyelination, as identified by manual pathologist annotation (**Fig. 6a**). This sample measures approximately 22.3 mm × 23.6 mm (567.7 mm^2^) and consists of 2,042,781 superpixels within the tissue, each measuring 8 µm × 8 µm. Similar to Sample 1, gene expression in Sample 2 was originally measured using six daughter captures from the 10x Visium platform, covering only a limited area of the tissue (**Supplementary Fig. 9b**). To demonstrate iSCALE’s out-of-sample prediction capability, we applied the model trained on Sample 1’s daughter captures to predict super-resolution gene expression and performed tissue segmentation in Sample 2. For comparison, we also applied iSCALE directly to the six daughter captures from Sample 2 for in-sample predictions. As shown in **Fig. 6a**, the segmentation obtained from out-of-sample predictions in Sample 2 resembles the in-sample segmentation achieved using Sample 2’s own daughter captures, and aligned well with the overall tissue structure annotated by the pathologist. For example, iSCALE’s out-of-sample segmentation accurately identified distinct anatomical regions, including the lesion area, normal appearing white matter, and cortex. We further validated iSCALE’s out-of-sample predictions using IHC staining for MOG and CD68 obtained from adjacent tissue sections. As shown in **Fig. 6b**, despite being trained on a different sample, iSCALE accurately inferred the expected locations for both markers.

**Fig. 6.**
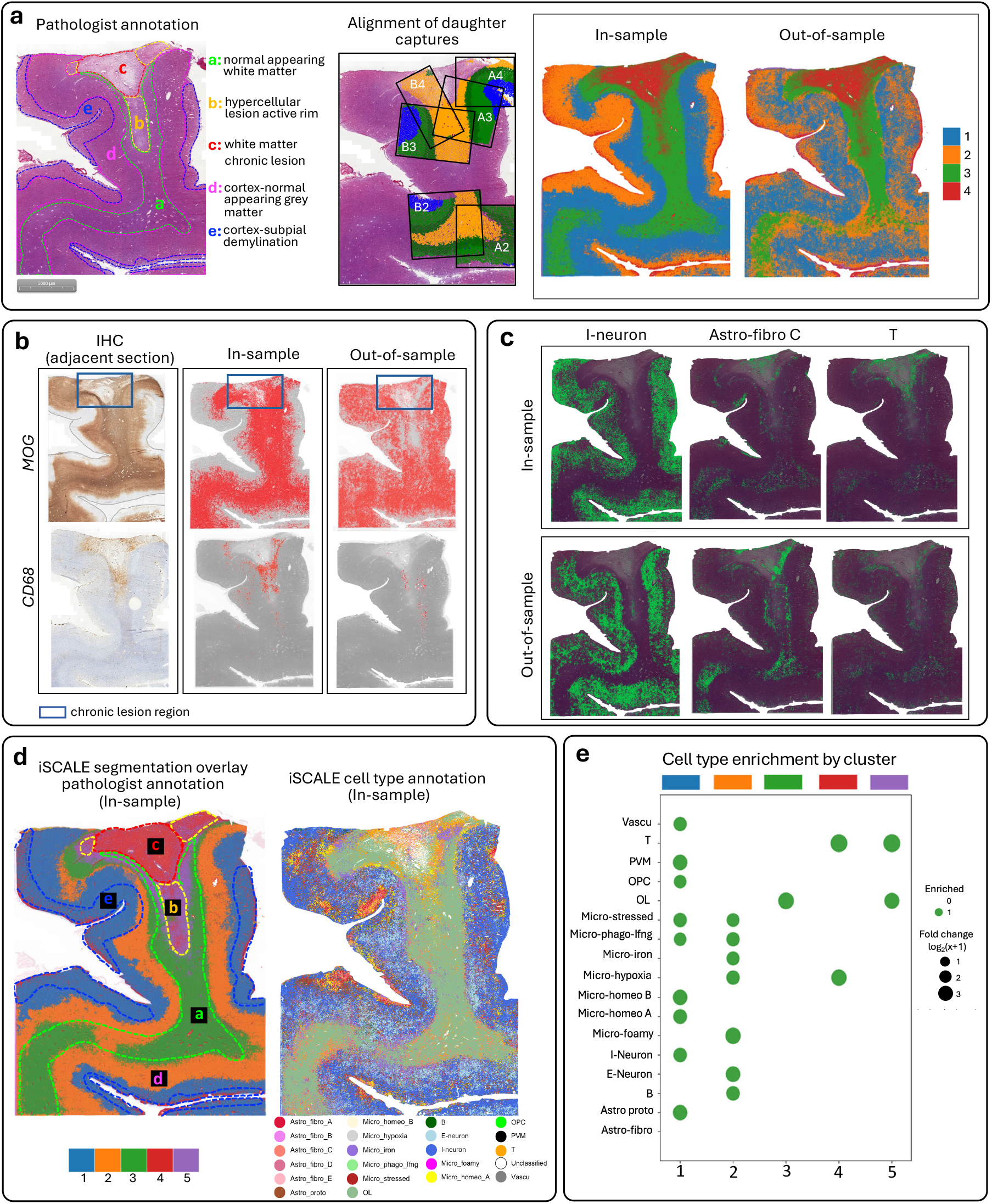
Application of iSCALE to an additional large postmortem human brain sample with MS. **a**, Comparison of iSCALE segmentations for in-sample and out-of-sample predictions for Sample 2. The panels include the mother H&E image with pathologist manual annotation, alignment of six daughter captures onto the mother image, and segmentations using iSCALE’s predicted gene expressions. In-sample prediction was based on a prediction model trained using the Visium data generated from the 6 daughter captures for Sample 2. Out-of-sample prediction was based on a prediction model trained using the Visium data generated from the 11 daughter captures for Sample 1. **b**, Experimental validation of iSCALE’s out-of-sample predictions by IHC staining, where the IHC stains were taken from adjacent tissue sections for critical markers including MOG and CD68. **c**, Comparison of cell type annotations for in-sample and out-of-sample predictions. Displayed are the annotations for inhibitory neurons (I-neuron), fibrous astrocytes subpopulation C (Astro-fibro C), and T cells. **d**, Left: Segmentation based on iSCALE’s in-sample predictions, overlaid with the pathologist’s annotations. Right: Cell type annotations derived from iSCALE’s in-sample predictions. **e**, Cell type enrichment analysis by cluster using iSCAEL’s in-sample predictions.

Next, we evaluated iSCALE’s performance in cell typing for out-of-sample predictions (**Supplementary Fig. 14**). This is more challenging than the benchmarking experiment, as the two normal samples, though from different patients, are structurally similar. In contrast, the two MS samples, while originating from the same brain, exhibit distinct lesion characteristics. Sample 1 contains a complete white matter inactive lesion core, whereas Sample 2 includes a chronic active lesion and areas of active and chronic subpial cortical demyelination. Despite these significant differences between Sample 1 and Sample 2, the out-of-sample predictions in Sample 2 successfully inferred inhibitory neurons within the grey matter, with the locations closely matching those inferred from in-sample predictions (**Fig. 6c**). In addition, the out-of-sample predictions in Sample 2 mapped T cells to the expected lesion rim, and astrocytes to the expected normal appearing white matter.

Although iSCALE’s out-of-sample prediction is currently limited to revealing broad tissue architecture through segmentation and identifying a limited number of cell types, it is worth noting that the two samples are different in their lesion characteristics. Importantly, iSCALE’s in-sample segmentation for Sample 2 closely aligned with pathologist’s manual annotation and detected the expected cortical layer structure when the number of clusters increased (**Supplementary Fig. 15**). We also observe strong alignment between the identified clusters and the iSCALE cell type annotation (**Fig.6d**). Notably, cluster 3 (green) corresponded well with oligodendrocytes and the normal appearing white matter in the pathologist annotation. Additionally, astrocytes (astro-fibro C), T cells, and oligodendrocytes were identified throughout cluster 4 (red) corresponding to the white matter chronic lesion (**Fig. 6d; Supplementary Fig. 16**). Furthermore, predicted biomarkers using in-sample predictions revealed tissue regions that posed challenges for manual annotation. For example, the subpial demyelination in the cortex was annotated by the pathologist using the mother H&E image and IHC staining for MOG obtained from an adjacent tissue section (**Fig. 6b; Supplementary Fig. 17**). This introduced uncertainty in defining the borders of subpial demyelination, highlighting the challenges pathologists face when relying on IHC staining from different adjacent slices. In contrast, iSCALE provides simultaneous predictions for all markers and enables automatic annotation on the same large tissue sample, offering a more integrated and consistent approach for tissue analysis. As shown in **Fig. 6e**, the absence of oligodendrocytes in the cell type enrichment analysis confirms that cluster 1 corresponds to the subpial demyelination region.

## Discussion

In summary, we have presented iSCALE, an innovative framework for rapid gene expression prediction, cell type annotation, and tissue segmentation for comprehensive ST analysis of large-sized tissues, extending beyond the capabilities of conventional ST platforms. By integrating high-resolution histology images with ST data from a small set of daughter captures, iSCALE effectively processes large tissue samples. The biology and physiology of diseased tissues involve a complex interplay of spatially varying gene expression and cellular heterogeneity in tissue architecture. Using benchmarking data from gastric cancer and normal gastric tissues, we demonstrated iSCALE’s effectiveness in semi-automatic daughter capture alignment and its ability to accurately predict super-resolution gene expression across the entire large tissue. When applied to MS brain samples, iSCALE further demonstrated its utility in assisting scientists in high throughput and unbiased tissue annotation of entire lesions, inferring cell type compositions, and pinpointing regions of interest with features not easily discernible through human visual assessment alone.

Although we demonstrated the utility of iSCALE primarily using Visium, the framework is versatile enough to be applied to other ST platforms^26–28^, and help improve our ability to uncover meaningful functions and insights from genomic data. For example, iSCALE can employ a set of selected field of views (FOVs) in CosMx to train the gene expression prediction model, which will enable the prediction of single-cell resolution gene expression in tissue regions outside those FOVs, thereby expanding gene expression coverage to the entire tissue section.

In both benchmarking and real-world applications, we demonstrated iSCALE’s effectiveness in aligning daughter captures. The gastric and brain tissue samples we analyzed have distinct tissue structures. However, we recognize that the semi-automatic alignment algorithm may encounter challenges in tissues lacking clear spatial patterns. Addressing this limitation is an area for future improvement.

This paper also highlights iSCALE’s ability to perform out-of-sample prediction—a highly challenging task due to the absence of observed gene expression and heterogeneity between the training and test data, especially in disease contexts. While our results remain preliminary, given the limited training data in both the benchmarking and real-world scenarios, they carry significant practical implications. H&E-stained histology images are widely accessible, cost-effective to generate, and well-suited for large tissue sections. We anticipate that as more ST data are available, iSCALE will benefit from the increased training data. iSCALE’s capability for out-of-sample prediction could substantially reduce experimental costs, making large-scale ST studies more feasible.

In summary, iSCALE is a computational tool that can help enhance our understanding of spatial tissue architecture at the cellular level, offering valuable insights for a wide range of human diseases. This capability has the potential to make significant contributions to future advancements in human healthcare. With the increasing adoption of spatial transcriptomics in biomedical research, we anticipate that iSCALE will have broad applications, particularly in studies involving large-sized tissues.

## Acknowledgments

M.L. was partly supported by NIH grants R01HG013185, R01LM014592, R01HL171595, and a grant from Biogen Inc.

## Author contributions

This study was conceived of and led by M.Li, B.Z., and T.H.H. A.S. designed the model and algorithm with input from M.Li and D.Z.. A.S. implemented the iSCALE software and led data analyses with input from all other coauthors. M.Loth developed the semi-automatic daughter capture alignment algorithm. M.Loth, C.L., S.Y., H.Y., S.P., W.H. participated in data analyses. T.H.H, J.H.P., J.-Y.S., J.R.C., I.J., M.K., and I.B. generated and processed the Xenium data for a gastric cancer sample and two normal gastric samples. J.H.P. annotated the gastric cancer and normal gastric tissue samples. E.P. and S.J.C. annotated the MS brain samples. H.-H.T., T.L.R., T.C., P.C. provided biological interpretation of the MS results. A.S., M.Li, and B.Z. wrote the paper with feedback from the other co-authors.

## Methods

### Semi-automatic alignment of daughter captures onto mother image

A critical step in iSCALE is aligning daughter captures onto the mother image. Given that the daughter captures are often derived from different tissue sections and exhibit drastically different shapes, existing alignment methods fall short of adequately addressing the complexity of this data. To overcome these challenges, we developed a semi-automatic, human-in-the-loop alignment approach. This method streamlines the alignment process while maintaining the flexibility needed to handle complex data structures. The approach involves the following steps:

**Step 1: Clustering spatial gene expression data in daughter captures**. Apply spatial clustering algorithms, e.g., BayesSpace^29^, to cluster gene expression data in daughter captures. We recommend generating 2 to 4 clusters to minimize noisy clustering patterns. The resulting clusters can then be used to facilitate the identification of key points in Step 2.

**Step 2: Identify key point pairs between a daughter capture and the mother image**. Identify at least 4 key point pairs between a daughter capture and the mother image. Use the clustering results from Step 1 as a guide to locate key points by comparing the clustering patterns observed in the daughter capture with the histological patterns in the mother image.

**Step 3: Find a transformation for alignment**. Use the key point pairs identified in Step 2 as input for the RANSAC algorithm implemented in OpenCV to determine a transformation between the daughter capture and the mother image. This transformation enables the alignment of the daughter capture onto the mother image.

**Step 4: Align a new daughter capture**. Select a new daughter capture and identify key point pairs between this capture and the mother image. Leverage the clustering results of both the new daughter capture and the previously aligned daughter capture to guide the identification of key point pairs between the new daughter capture and the mother image. Apply the RANSAC algorithm, as outlined in Step 3, to find the transformation, and align the new daughter capture onto the mother image.

**Step 5: Repeat alignment of remaining daughter captures**. Continue the above alignment process until no additional daughter captures can be aligned.

The semi-automatic procedure described above requires minimal manual work and is relatively straightforward to implement. **Supplementary Fig. 2** provides an example to illustrate how this alignment algorithm works.

### BayesSpace clustering of gene expression in daughter captures

For data analyzed in this paper, the initial clustering of the daughter captures was performed using BayesSpace^29^. In this analysis, we combined gene expression data across all daughter captures but shifted their spatial locations to prevent overlap. This allowed BayesSpace to utilize spatial information within each sample while ensuring that cluster assignments remained consistent across the samples.

### 10x Xenium experiment of human gastric tissue specimens

A gastric cancer sample was previously analyzed using the 10x Xenium platform, as described in Coleman et al.^30^. In this study, we generated two additional 10x Xenium datasets from normal gastric tissue samples. Formalin-fixed, paraffin-embedded sections of two normal gastric tissue blocks were cut with 5 μm thickness into corresponding Xenium® slides. Deparaffinization was performed by incubating slides at 60°C for 2 hours, followed by 10 minutes in xylene jar 1 and 3 minutes in xylene jar 2, 100% ethanol jar 1, 100% ethanol jar 2, 96% ethanol jar 1, 96% ethanol jar 2, and 70% ethanol, respectively. After a 20 second water rinse, RNA decrosslinking was performed at 80°C for 30 minutes. The 10x Genomics Human Multi-Tissue and Cancer gene panel V1 was used to hybridize probes for 377 genes for 18 hours at 50°C. Following 10x Genomics protocol CG000582, revision E, unhybridized probes were washed away at 37°C for 30 minutes, followed by probe ligation for 2 hours at 37 °C, and rolling circle amplification at 30°C for 2 hours. After washing, background autofluorescence was quenched, and nuclei where fluorescently stained. Prepared tissue sections were then imaged using the Xenium® Analyzer instrument following 10x Genomics protocol CG000584, revision C. The instrument automatically performed fluorescent probe hybridization, imaging, and probe removal as a series of 15 cycles using firmware version 1.7.6.0. The Xenium® Analyzer then automatically processed and analyzed the images by performing image co-registration, probe quality filtering, nuclei and cell segmentation using software analysis version xenium-1.7.1.0.

### Tissue sectioning and immunohistochemistry of postmortem human brain tissue specimens

Postmortem human brain tissues were obtained from a patient with secondary progressive multiple sclerosis. Fresh frozen samples were obtained from the Imperial College London. Frozen human tissue blocks were allowed to equilibrate to −23°C inside the cryostat for at least 30 minutes using a minimal amount of OCT for mounting. Blocks were gently faced to allow collection of full-face sections. Prior and following collection of tissue samples on Visium slides, four 10 μm-thick sections were collected in series onto individual Plus Gold glass slides and were fixed by incubation in 10% NBF for 15 minutes followed by two 10-minute washes in PBS. Glass slides were then removed from PBS and allowed to air dry for standard H&E, CD68 and MOG staining. CD68 and MOG staining were carried out as follows on a Leica Bond RXm or Ventana DISCOVERY Ultra automated stainer using standard chromogenic methods. For antigen retrieval (HIER) and permeabilization, slides were heated in a pH=9 EDTA-based buffer for 25m at 94°C, followed by a 30-minute antibody incubation (CD68 1:2250 Cell Signaling rabbit clone D4B9C, 76437 and MOG 1:2000, Abcam rabbit clone EPR22629-310, ab233549). Bound antibodies were detected with an HRP-conjugated, anti-rabbit secondary polymer, followed by chromogenic visualization with diaminobenzidine (DAB) and hematoxylin counterstaining. Oil red O (ORO) staining, previously performed and provided by Imperial College London (ICL), was used to identify lipid laden microglia/macrophage in this study. Whole slide images were acquired on a Pannoramic 250 digital scanner (PerkinElmer).

### Characterizing white matter, grey matter demyelinated lesion and active phase RIM in the MS brain samples

MS lesion types were determined by the relative absence of MOG immunoreactivity indicating demyelinating/demyelinated areas using MOG. The reactivity of lesion types was further characterized by the activity of CD68 and ORO stain. White matter lesion types were classified into active, chronic active, and chronic lesion according to Kuhlmann *et al*. ^31^. Briefly, white matter active lesion (WMAL) was characterized by the loss of MOG reactivity and the presence of large numbers of lipid laden macrophage/microglia cells with CD68 reactivity and positive ORO stain. White matter chronic active lesion (WMCAL) was characterized by demyelinated lesion with CD68 immunoreactivity form a rim at the lesion border. The active RIM was annotated surrounding WMCAL with the presence of an extensive CD68 positive cellular infiltrate, corresponding to macrophages/activated microglia as described by Absinta *et al*. ^32^. Inactive lesion was annotated at demyelinated lesion with a few ramified CD68 positive cells. Cortical grey matter lesion was characterized by the loss of MOG reactivity at grey matter regions and further subtyping into subpial, leukocortical and intracortical lesion^33, 34^. Briefly, lesions showing loss of myelin indicated by MOG staining from the outer layer into the cortex were classified as subpial lesion. Leukocortical lesion is characterized by the demyelination that is present at the cortical subcortical border that affects the grey as well as the white matter. Whereas intracortical lesion is characterized by grey matter demyelination located around a blood vessel.

### 10x Visium experiment in the MS brain samples

To collect tissues onto Visium Spatial Gene Expression slides (10X Genomics Slide Kit, Ref No. 1000185), a pre-chilled razor blade was used to gently score the surface of the frozen tissue block in a grid format to subdivide the tissue for placement on individual capture areas. Slides were pre-chilled prior to collection of tissue sections and individual sections were collected onto slides by placing a fingertip on the back of the slide to promote tissue capture. Frozen slides were then processed for spatial transcriptomic analysis according to the manufacturer’s instructions (10X Visium “Tissue Staining & Imaging.” Sections were heated at 37C for 1 minute, followed by methanol fixation at –20C for 30 minutes. Samples were then stained with hematoxylin and eosin. Brightfield images were collected on an Olympus VS120 Virtual Slide Microscope equipped with a VC50 camera using a UPLSAPO 20X objective. After imaging, tissue permeabilization was carried out using the supplied Permeabilization Buffer at 37C for 6 minutes. Permeabilization time was selected based on a prior optimization experiment in mouse tissues using Visium Spatial Tissue Optimization slides (10X Genomics, Ref. No. 1000193). Following permeabilization, reverse transcription was carried out on-slide according to Visium guidelines.

### Integrating multiple ST captures to construct iSCALE gene expression training data

Integrating the location and spot-level gene expression information from multiple Visium captures involves combining information from individual ST experiments to construct the iSCALE training dataset. To construct the iSCALE training dataset for the gene expression features, we strategically combined information from individual ST experiments by integrating the location and spot-level gene expression information from multiple daughter captures. Let *ϕ*_1_, *ϕ*_2_,…, *ϕ*_*s*_ denote the spatial coordinates (*X*_*s*_, *Y*_*s*_) for each daughter capture, where *S* is the total number of ST daughter captures used for training the iSCALE model. We denote the number of spatial observations in daughter capture *s* to be *q*_*s*_, where 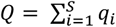 is the total number of observations in the joined training set. To combine the spatial locations across *S* daughter captures into a unified dataset *L* ∈ ℝ^2^, we concatenate the sets of aligned coordinates such that *L* = *ϕ*_1_ ∪ *ϕ*_2_ ∪ … ∪ *ϕ*_*S*_, where *L* is of size *Q* × 2.

Next, we combine gene expression data from multiple daughter captures. Let *g*_*s*_ denote a *q*_*s*_ × *w*_*s*_ sized matrix containing gene expression measures for daughter capture *s*, where each row corresponds to a spatial measurement (i.e. spot or cell) and each column represents a gene. Entry *g*_*sij*_ in the matrix denotes the expression level of gene *j* at spatial location *i* in capture *s*. Therefore, we have matrices *G*_1_, *G*_2_, …, *G*_*S*_, as the gene expression input into the iSCALE pipeline. We define *τ*_*s*_ as the set of all genes measured in daughter capture *s* with length *w*_*s*_, and let *γ* denote the set of common genes between all *S* daughter captures with length |*γ*| = *φ*. Thus, *γ* = *τ*_1_ ∩ *τ*_2_ ∩ … ∩ *τ*_*S*_ is defined as the intersection of all gene sets corresponding to each daughter capture in the training set. Next, daughter captures are subset to include measures for genes only within *γ*. Denote 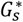 as the reduced daughter captures such that 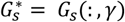 with size *q*_*s*_ × *φ* for daughter captures *s* = 1, …, *S*. Then, to integrate these compatible gene expression profiles into a single training dataset, denoted as *G*, we concatenate 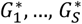 along the spatial axis such that 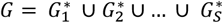 with dimensions *Q* × *φ*. Thus, the unified matrix, *G*, is used as a large training dataset encompassing partial regions of the entire tissue section analyzed by multiple ST captures. Additionally, iSCALE includes an integration step among captures from overlapping adjacent slices which involves smoothing based on dense neighborhoods. These neighborhoods are defined using a distance threshold *d* between spots, enabling the incorporation of information from adjacent slices. iSCALE then integrates the gene expression counts and spatial locations across dense neighborhoods forming the final expression matrix 𝔊. This approach ensures continuity and consistency across slices by effectively integrating spatially overlapping regions.

After forming the gene-expression training dataset, iSCALE automatically identifies the top set of highly variable genes (HVGs) across all spatial locations in the training set. The user inputs the desired number of HVGs, for iSCALE to select. Additionally, users have the flexibility to easily modify the gene list to include additional genes of biological or clinical interest for their study. Let ℧ denote the final set of genes used as input to train the iSCALE model, where ℧ ⊆ *γ* and the total number of genes used for model training is *K, K* = |℧|. Thus, we subset 𝔊 such that 𝔊^*^ = 𝔊 (:, ℧).

### Normalization of gene expression among daughter captures to mitigate batch effects

Integrating gene expression data from multiple ST captures can pose the challenge of batch effect. Consequently, this may obscure genuine biological signals and impede data integration for the training dataset. As illustrated in **Supplementary Fig. 10** (left panel), the integrated gene expression data reveals a prominent batch effect for the MS Sample 1 analysis, characterized by distinct groupings of daughter captures visible within the UMAP space. The iSCALE preprocessing pipeline automatically performs normalization of gene expression data to be scaled within the range of 0 and 1 for each gene across all daughter captures. This is achieved using a min-max scaling approach. Let 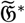 denote the normalized batch corrected joint gene expression data set, where 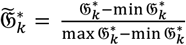, for each gene *k* ∈ {1, …, *K*} in the joint training set where 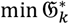 and 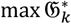 represent the minimum and maximum expression value of gene *k* across all spots in the integrated dataset. As shown in **Supplementary Fig. 10** (right panel), this scaling approach effectively removed the batch effect observed in the MS study data, resulting in the mixture of expression profiles from spots across different captures. Furthermore, following preprocessing, iSCALE automatically generates two UMAP plots, each colored by ST capture, to visualize the integrated training dataset before and after normalization. This feature enhances visualization for users, enabling them to observe the extent of batch effects within the integrated gene expression dataset and assess whether the automatic normalization approach in the iSCALE pipeline effectively mitigates the issue. If there appears to still be concerns in these projection visualizations, we recommmend considering alternative batch effect correction methods such as Z-score normalization or utilizing specialized batch correction packages such as CarDEC^35^ or Scanarama^36^ before running the iSCALE prediction model.

### Extraction of histology image features

Although histology images have been leveraged to analyze ST data in previous works^11, 16, 37, 38^, their use of histology information does not fully leverage the rich cellular information provided by high-resolution histology images. In practice, a pathologist examines a histology image in a hierarchical manner. In this process, the first step is to identify a region of interest (ROI) through the examination of high-level image features that capture the global tissue structure. After a ROI is identified, low-level image features that reflect the local cellular structure of the tissue are examined. To mimic this process computationally, we recently developed a hierarchical image feature extraction approach that captures both global and local tissue structures. This histology image feature extraction approach employs the Vision Transformer (ViT)^39^ in a hierarchical manner and extracts essential tissue details by breaking down the tissue image processing into stages. It starts by analyzing smaller tissue sections for the fine details, subsequently scaling up to grasp broader tissue patterns. Details of this image feature extraction were described in iStar^14^.

### Prediction of super-resolution gene expression in large tissue samples

Previously, the iStar^8^ method was developed for the purpose of enhancing the spatial resolution of a given standard sized ST dataset, where the training sample is the ST sample itself. For the problem considered here, there are multiple ST samples obtained from different daughter captures, and large tissue regions with no gene expression data measured in the target mother image. To utilize such data, we build upon the iStar method to incorporate multiple training samples and perform out-of-sample super-resolution gene expression prediction for the missing tissue regions. Recall, 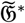 is the batch corrected normalized genes expression matrix containing the integrated measures from *S* daughter captures across *K* genes. We performed feature extraction on the large-scale H&E image to obtain 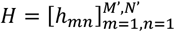 and weakly supervised gene expression prediction by training on the combined daughter data, 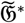. From there, we were able to perform cellular level super-resolution gene expression prediction, whole capture segmentation, and infer the cell type environment of the tissue. The method is capable of accurately predicting gene expression patterns and spatial relationships across the entire tissue section, even in regions not directly measured by Visium captures. The iSCALE model integrates information from multiple ST captures and performs out-of-sample prediction for regions of large tissue containing no gene expression information by leveraging information learned between the extracted histology features and the integrated ST dataset using an extension of the iStar spot-level loss function^8^ and weakly supervised learning model to integrate information from multiple ST captures when training. To express the loss function, recall *Q* denotes the total number of spatial observation measured across *S* daughter samples, *k* represents the number of genes to be predicted, *g*_*k*_ is the gene expression prediction model for gene *k*, 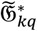 is the observed gene expression for gene *k* at spot *q* in the integrated and batch corrected training set across *S* ST samples, ℳ_*q*_ is the spot mask of *q* (that is, the collection of superpixels covered by spot *q*), and *h*_*mn*_ be the histology feature vector at superpixel (*m, n*). Then the weakly supervised loss function is

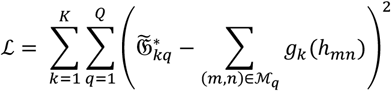

After model training, the predicted gene expression for gene *k* at superpixel (*m, n*) is represented as 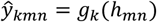, resulting in the gene expression image 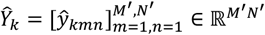 for the large scale mother image.

### Tissue segmentation

The predicted gene expression can be used for tissue segmentation through clustering analysis with the k-means algorithm. iSCALE provides two options for this clustering analysis. The first option uses the last embedding layer from the neural network employed for gene expression prediction as input, while the second option uses principal components derived from the predicted gene expression as input. By the end of the clustering analysis, iSCALE partitions the tissue into functionally distinct regions in an unsupervised manner based on their gene expression profiles.

#### Inferring cell type composition of large tissues using a percentile score-based approach

To aid in assigning meaningful interpretations to the segmented tissue regions, and provide insights into the composition and heterogeneity of cell populations, we perform cell type inference at the superpixel level across the entire tissue section. In this process, each superpixel is treated as an artificial cell, and its cell type is inferred using the iSCALE predicted gene expressions alongside a marker gene reference panel. The original cell typing approach implemented in iStar often produces over-dominant cell type predictions, where a single cell type is predicted across the entire tissue section. This issue can arise when the marker list includes genes with outlier expression levels or overestimated expression measures. Therefore, iSCALE introduces several key modifications to ensure more robust cell-type annotation.

Let *V* be the total number of candidate cell types in the marker reference set. For each cell type *v* ∈ {1, …, *V*}, suppose we have a list of marker genes names *ψ*_*v*_. In iStar, for each marker gene *k* ∈ *ψ*_*v*_, the predicted super-resolution gene expression image 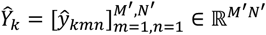 was standardized into the range of [0.0, 1.0] to obtain 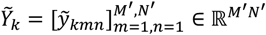, where 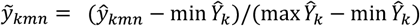, then the average marker score for each cell type was computed directly using 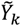.

In iSCALE, we first employ a filtering criterion based on the model’s prediction uncertainty for genes in the marker set. Our rationale is that if a marker gene cannot be predicted well in the training set, then this gene should be eliminated for cell typing. To do so, we compute the training RMSE for each gene in the model. We obtain a refined marker gene set 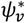 by removing all genes whose RMSE exceed the 95th percentile of the training RMSE for all genes as potential candidates from the cell-type marker set. Next, we iterate through each gene in the refined marker set, 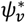, specific to each cell type and compute the percentile of each measure across all superpixels in the tissue. For each marker gene, the predicted super-resolution gene expression is transformed to 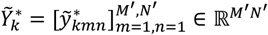, where we define 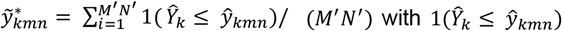 with 1(Ŷ_*k*_ ≤ Ŷ _*kmn*_) representing an indicator function that equals 1 if Ŷ _*k*_ ≤ Ŷ _*kmn*_ and 0 otherwise. This transformation still standardizes the data into the range of [0.0, 1.0], but removes the large impact of outlying or over predicted expression on the following calculation of the marker score. For each superpixel (*m, n*), we then compute the score for cell type *v* by averaging the percentile transformed gene expression for all its marker genes: 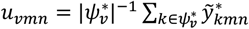 where 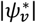 is the number of genes in 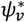. To infer the cell type of superpixel (*m, n*), let 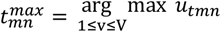 be the cell type with the maximal score and 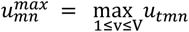 be the score of this cell type. Given a predetermined threshold *u*_*threshold*_ ∈ [0.0,1.0], if 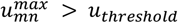, then the cell-type of superpixel (*m, n*) is predicted to be 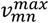; otherwise, the cell type of this superpixel is unclassified. In our experiments, we set *u*_*threshold*_ = 0.05 and found it effective in most cases.

The adjusted score-based approach allows the set threshold to be more consistent across different datasets that show varying magnitudes of imputed expression. It also lessens the somewhat arbitrary selection of thresholds commonly associated with the standard score-based method, providing a more interpretable alternative. Furthermore, the percentile adjustment helps mitigate the influence of outlier genes with high predicted expression levels, which could otherwise dominate the score function and lead to a single dominating cell-type label and is more robust to changes in the selected marker gene set. In additional to the percentile score-based approach that is used in our pipeline, the output from iSCALE allows users the flexibility to use any cell-type annotation method of their choice for the predicted super-resolution gene expression.

After inferring cell types, iSCALE can perform cell type enrichment analysis using the procedure described in the iStar^14^ paper. This involves examining each cell type-tissue cluster pair to identify biological activities within each cluster by determining which cell types are over-represented in the cluster.

### Evaluation metrics for gene expression prediction

For in-sample predictions, we calculated three metrics for each predicted gene. The first metric is root-mean-square error (RMSE), calculated as 1/√*n* of Euclidean distance between observed and predicted superpixel expression across *n* superpixels. The gene expression values were normalized to the range [0,1], making the RMSE a quick and direct metric to evaluate the accuracy of vectorized outcomes. The second metric is Structural Similarity Index Measure (SSIM), which assesses the similarity between two images, considering both intensity and spatial structures within the image. A higher SSIM score signifies greater similarity between two images, with a value of 1 indicating two images being identical. The third metric is Pearson Correlation between the observed and predicted expression values across all superpixels.

For out-of-sample predictions, where the range of gene expression in the test sample is unobserved, metrics that rely on gene expression magnitude are no longer suitable. Instead, we use Spearman Correlation, which evaluates prediction accuracy based on the ranks of gene expression. Additionally, we introduce a Chi-squared statistic to assess the concordance between the binarized gene expression patterns of the ground truth and predictions. In this analysis, gene expression is categorized as ‘expressed’ (above a pre-specified threshold) or ‘not expressed,’ allowing for a comparison of overall expression patterns. This Chi-squared statistic follows a χ^2^ distribution with 1 degree of freedom under the null hypothesis of no concordance between the predicted and the ground truth gene expression patterns. We calculate p-value based on this null distribution and adjust for multiple testing across all genes using Bonferroni correction.

Complete information about the notation described in the Methods section is available in

**Supplementary Table 2**.

### Neural network architecture and hyperparameter tuning

We trained a feed-forward neural network with four hidden layers, each containing 256 neurons and employing a leaky ReLU activation function (*α*=0.1). The model was trained with batch size of 100 using the Adam optimizer with a learning rate of 1e-3 for 1,000 epochs across three random initialization states. The loss function minimized the difference between predicted and observed gene expression, either at the spot level (iSCALE-Seq) or super-pixel level (iSCALE-Img), depending on the training data resolution. While no formal hyperparameter optimization was conducted, the architecture and training parameters were selected based on prior experience and performance evaluation using RMSE loss. Training and validation were performed on an NVIDIA GPU with 64 GB of memory to accommodate the largest sample.

### Computational efficiency

We ran iSCALE on a server equipped with an NVIDIA GPU (driver version 550.54.15, CUDA 12.4). For the MS data, we utilized a single GPU with 64 GB of memory for Sample 1 and 32 GB for Sample 2. The process of extracting histology image features, smoothing, and exporting data took approximately 13 minutes for Sample 1 and 15 minutes for Sample 2. Integrating data across daughter captures, training the gene expression prediction model (1,000 epochs, repeated for three random states), predicting gene expression, and exporting super-resolution gene expression data for 1,000 genes required 1 hour and 35 minutes for Sample 1 and 54 minutes for Sample 2.

## Data availability

The data analyzed in this paper will be made publicly available once the paper is accepted for publication.

## Code availability

The iSCALE algorithm was implemented in Python. The codes are publicly available on GitHub at https://github.com/amesch441/iSCALE.

