## Supplementary Information for "Scaling up spatial transcriptomics for large-sized tissues: uncovering cellular-level tissue architecture beyond conventional platforms with iSCALE"

**Supplementary Fig. 1** Study designs for spatial transcriptomics in large-sized tissue. **(A)** **Grid layout design:** In this approach, the tissue is divided into a grid, and each section (referred to as a "daughter capture") is measured using a platform like Visium. This layout allows for systematic spatial coverage of the tissue. **(B)** **Adjacent tissue slice design:** This design involves taking daughter captures from adjacent tissue slices. However, due to the challenges of tissue processing, the resulting daughter captures from different slices may only partially overlap in the X-Y plane. Additionally, some areas of the original tissue ("mother capture") may remain uncovered, leading to unmeasured gaps in the tissue data.

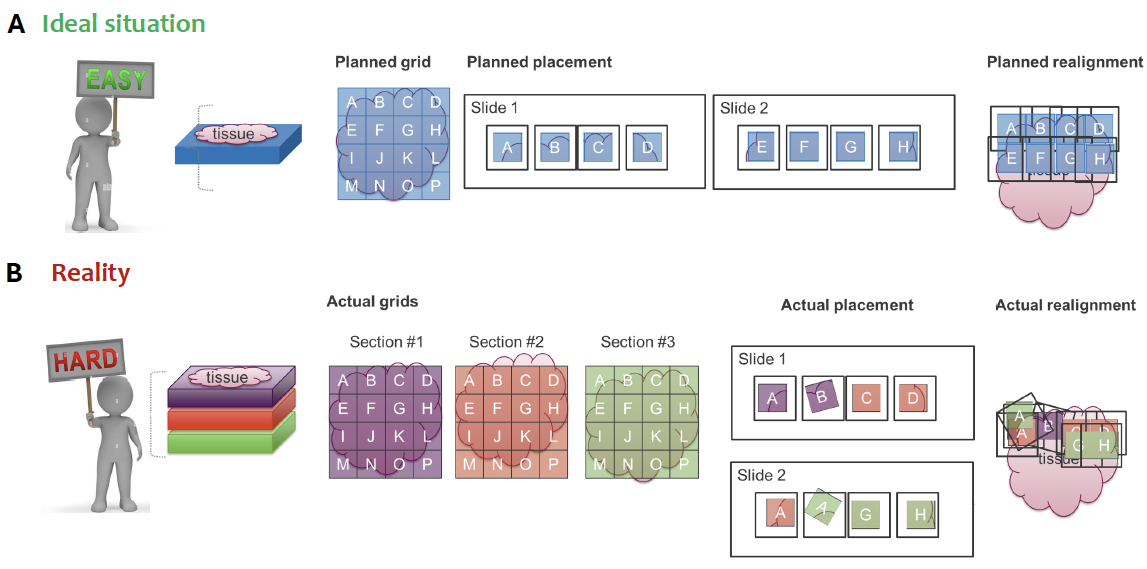

**Supplementary Fig. 2: Semi-automatic alignment method.** **(A)** Key point pairs plotted on MS Sample 1 mother image and Visium daughter capture 612-1 where Visium spots are colored by their respective BayesSapce clusters. These points were used to calculate the transformation needed to align one capture to the coordinate space of the mother image. **(B)** Aligned daughter capture (612-1) after the transformation. The alignment shows nice continuity in the spatial patterns between the cluster boundary and mother tissue boundary. **(C)** The final alignment of 11 daughter captures onto the mother image of Sample 1. **(D)** The final alignment of 6 daughter captures onto the mother image of Sample 2.

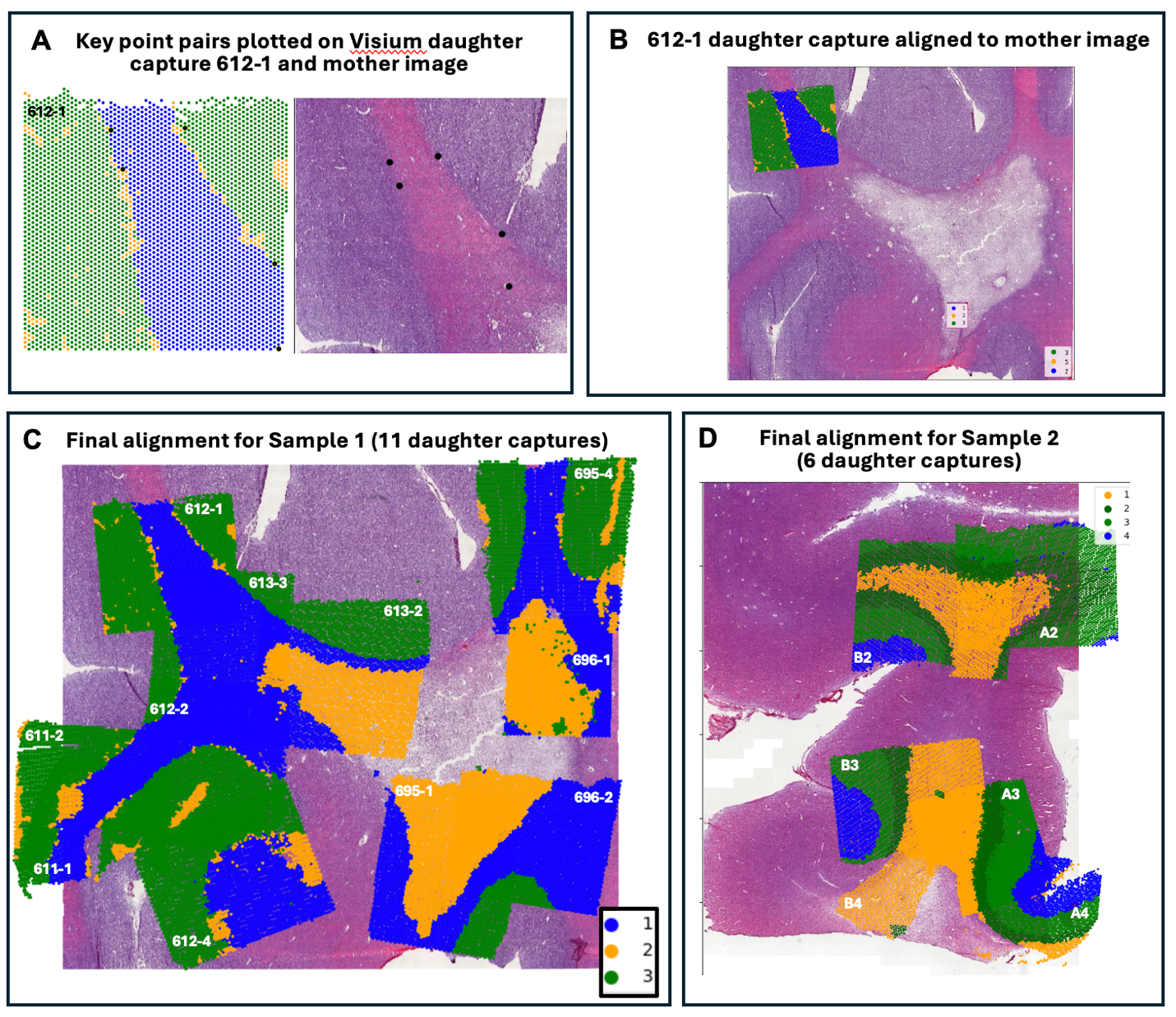

**Supplementary Fig. 3:** Visualization of the true cell locations across the gastric cancer tissue compared to the cell locations detected by RedeHist.

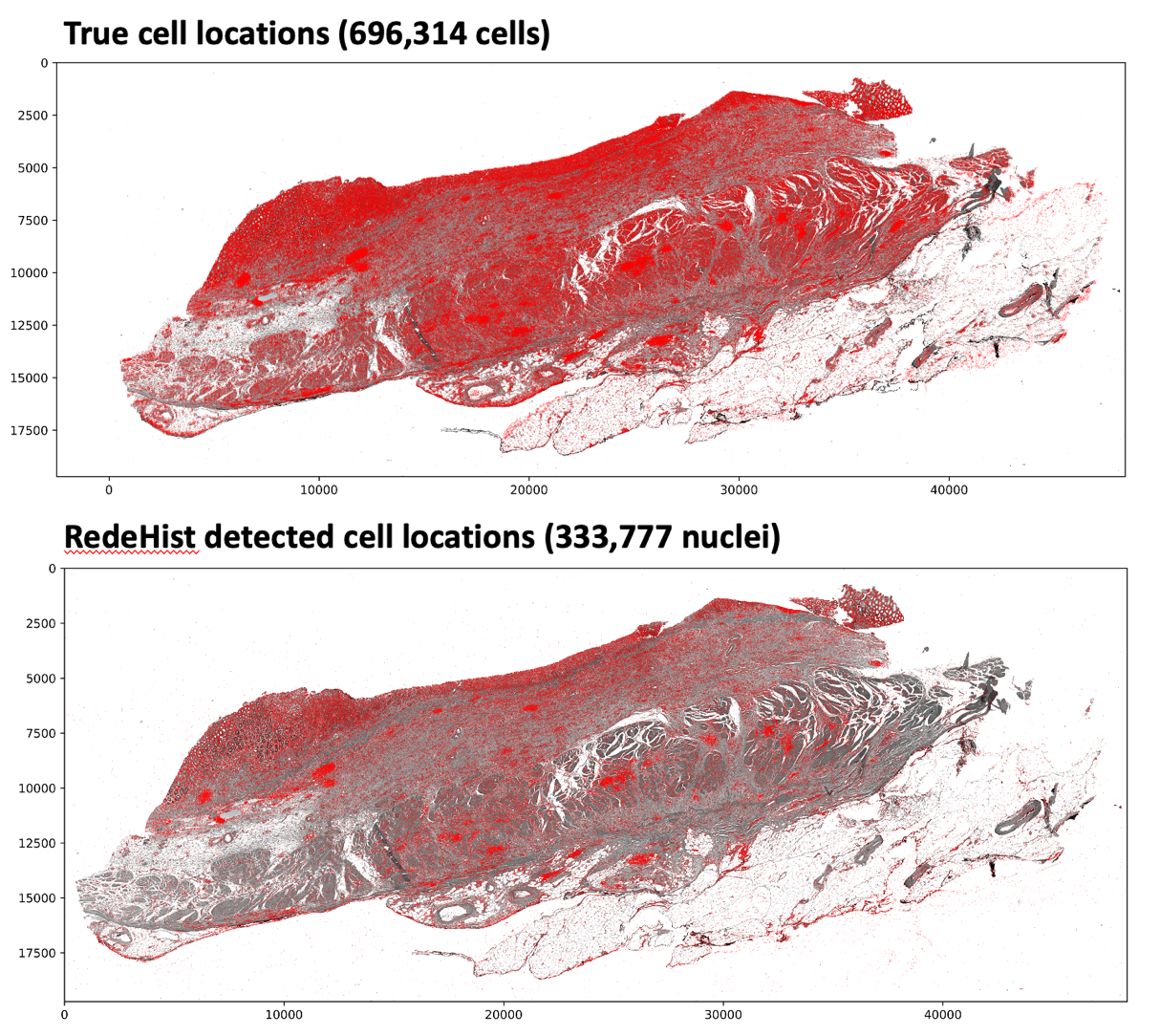

**Supplementary Fig. 4:** Gastric tumor border and intestinal metaplasia segmentation in the gastric cancer benchmarking data by iStar and RedeHist.

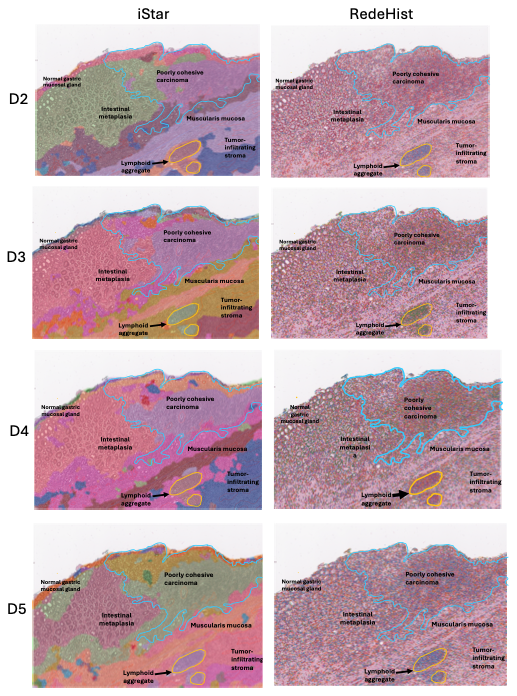

**Supplementary Fig. 5:** TLS detection in the gastric cancer benchmarking data by iStar and RedeHist.

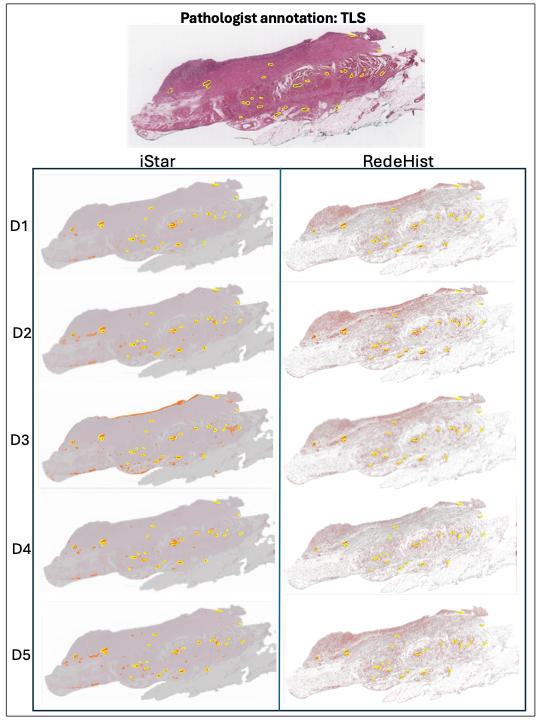

**Supplementary Fig. 6.** Visualization of additional genes predicted by iSCALE, iStar, and RedeHist in the gastric cancer benchmarking dataset. Since RedeHist relies on an external scRNA-seq reference, based on the reference dataset we used (Cheng et al., Gastroenterology. 2024 Dec;167(7):1345-1357.), RedeHist was only able to predict 38 of the top 100 highly variable genes. Visualization for additional genes can be found in

<https://upenn.app.box.com/s/ctpdj52xj4e5afvltvaoqsl7cinuekhq>

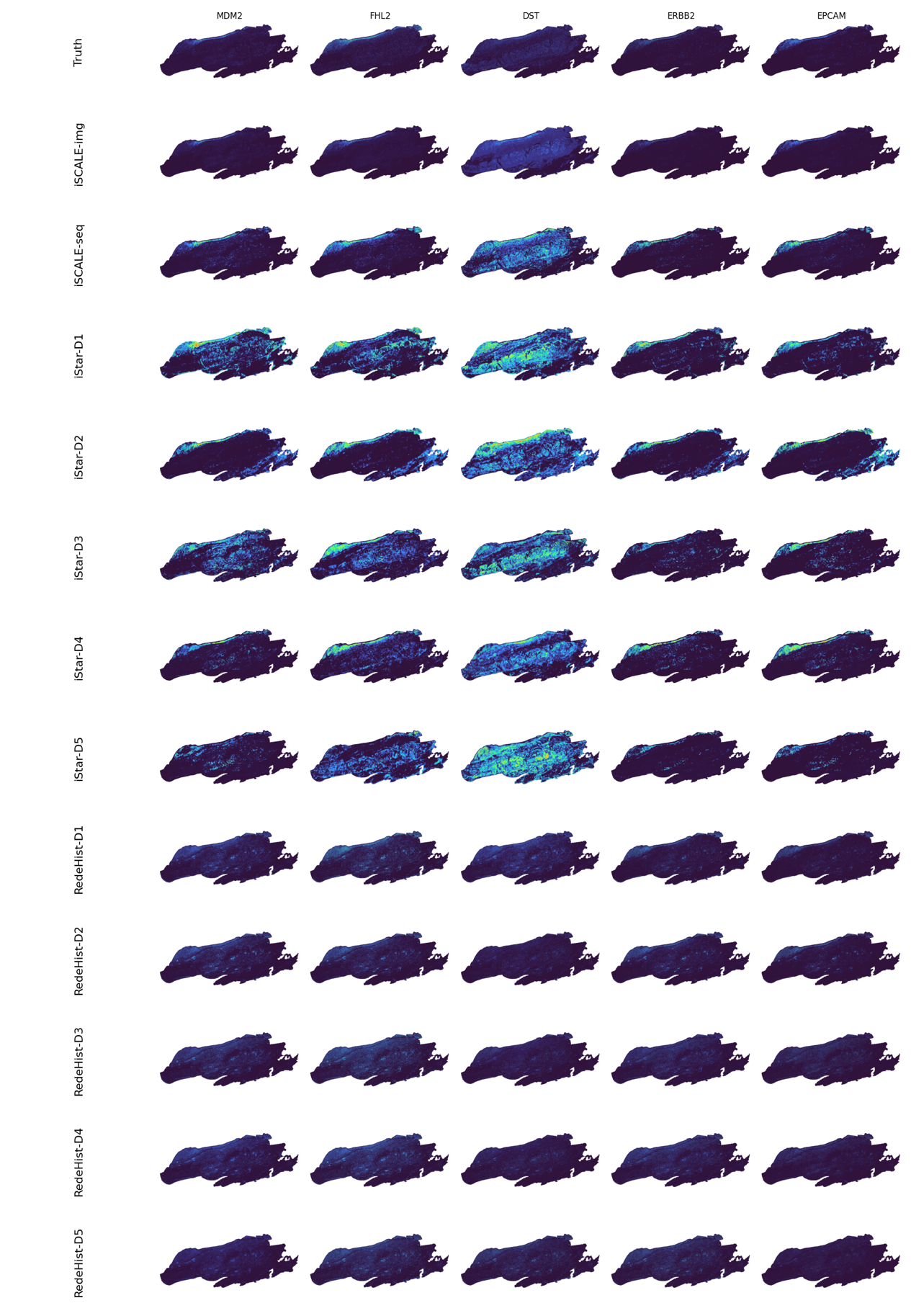

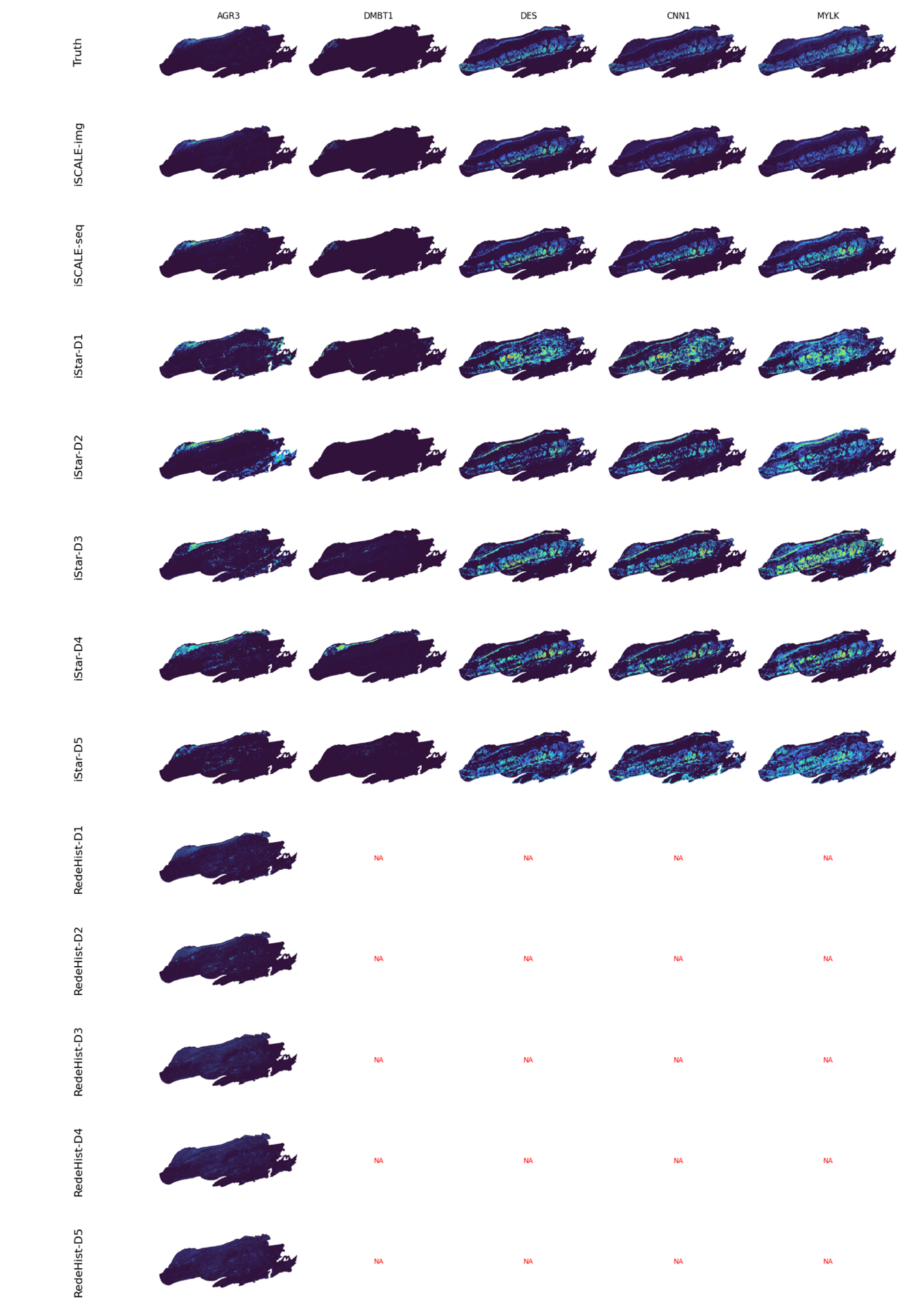

**Supplementary Fig. 7:** iSCALE’s semi-automatic alignment results for the benchmarking gastric tumor sample. The black box indicates the true location of each daughter capture, and the red box indicates the aligned location of the daughter capture. For each daughter capture, the number indicates the percentage of pixels that overlap between the aligned position and the true position.

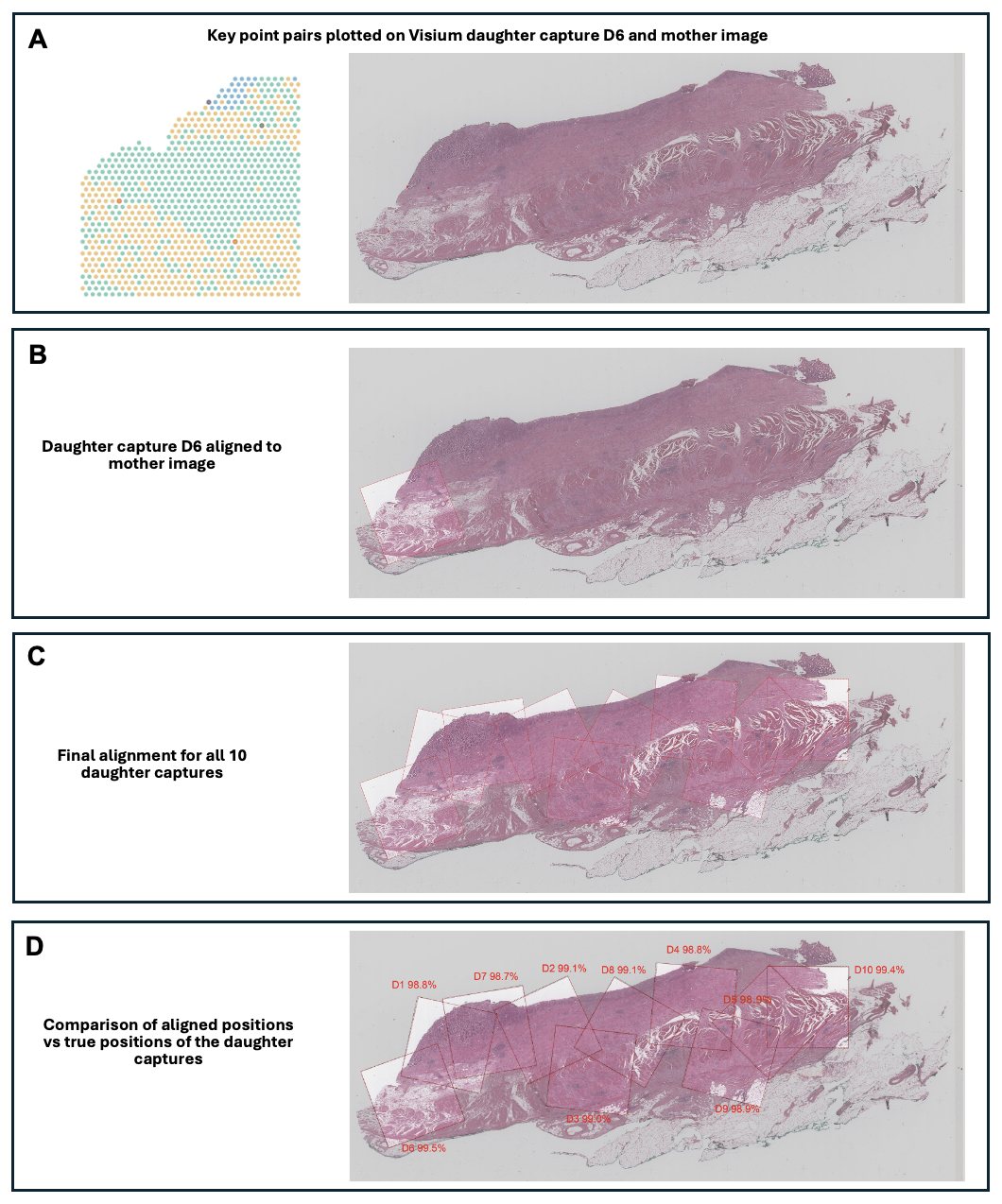

**Supplementary Fig. 8:** Visualization of additional genes predicted by iSCALE in-sample vs out-of-sample prediction in Normal 2 of the normal gastric tissue benchmarking data. Visualization for additional genes can be found in <https://upenn.app.box.com/s/ctpdj52xj4e5afvltvaoqsl7cinuekhq>

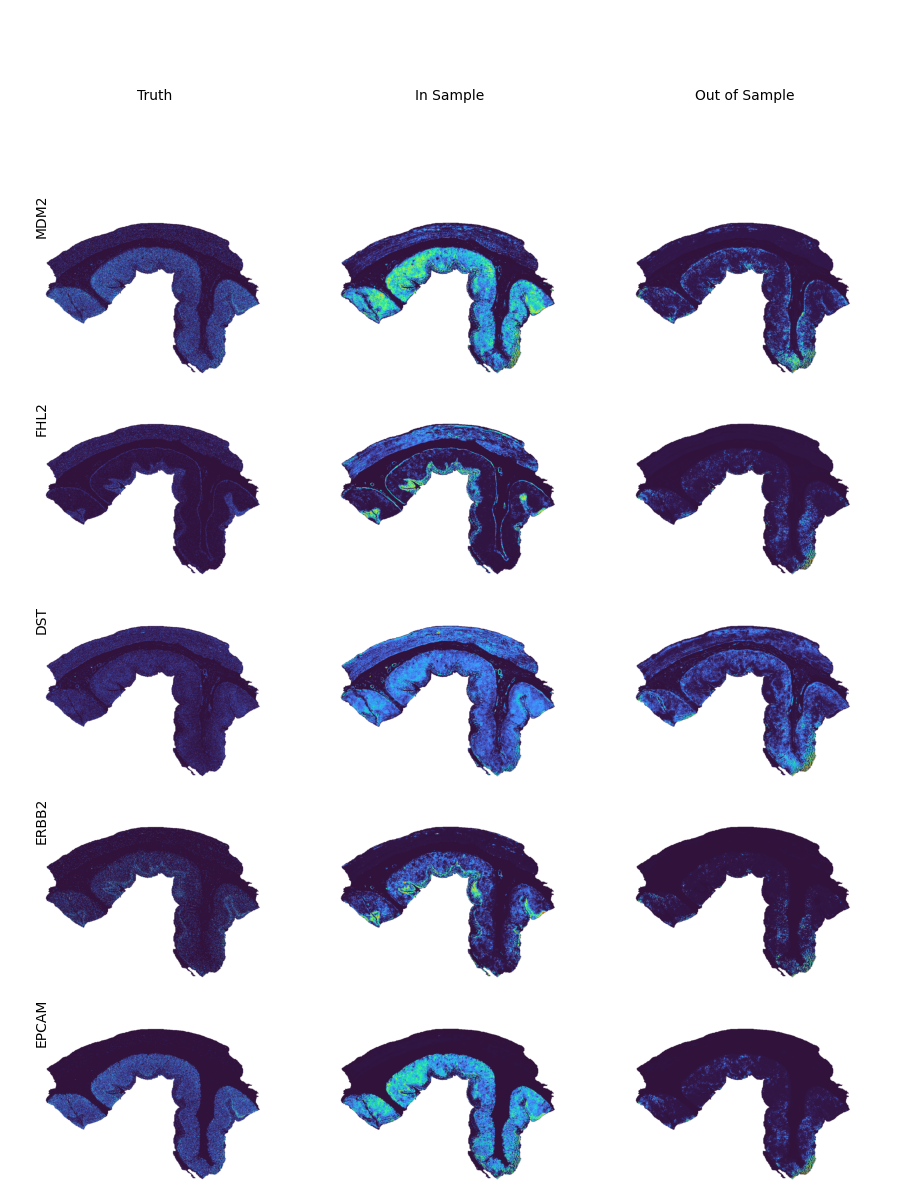

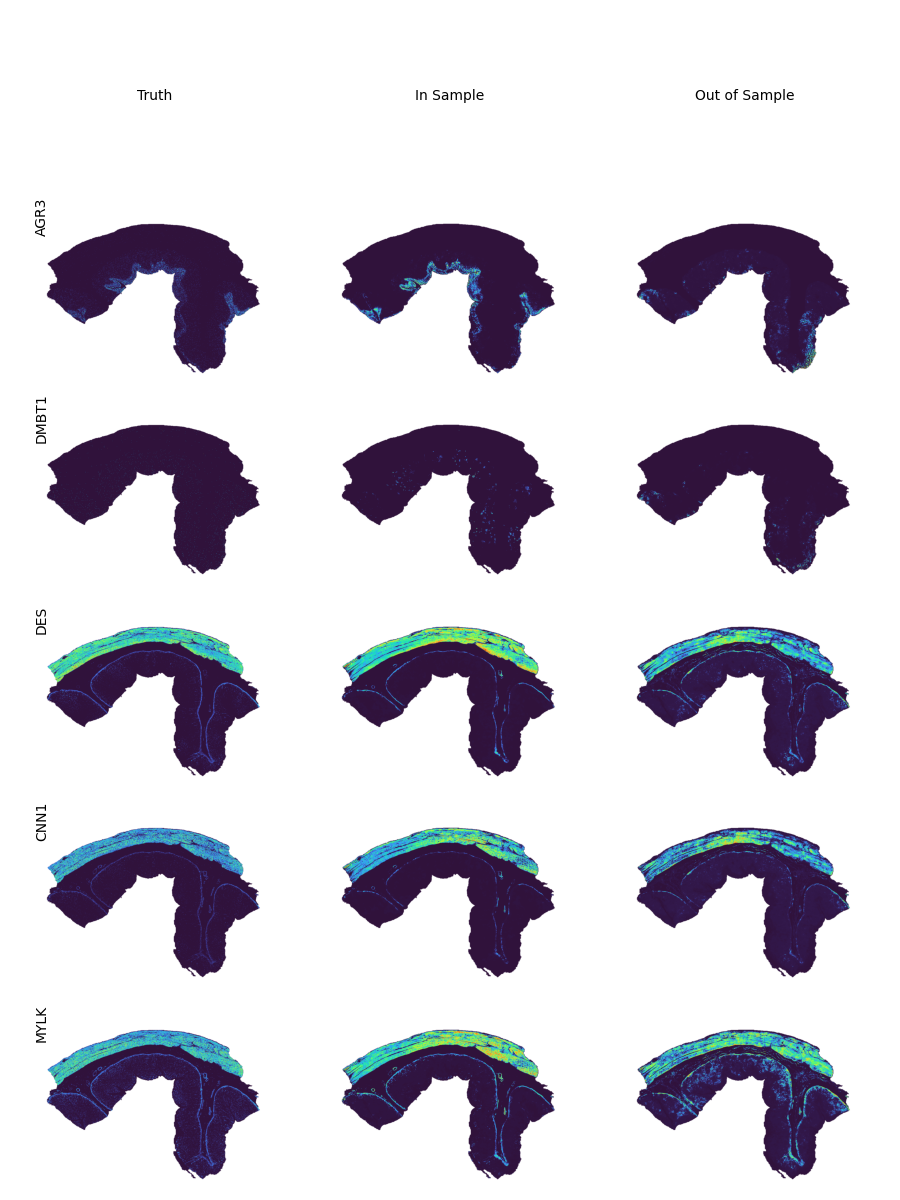

**Supplementary Fig. 9:** Study design and Visium data generation in the multiple sclerosis study.

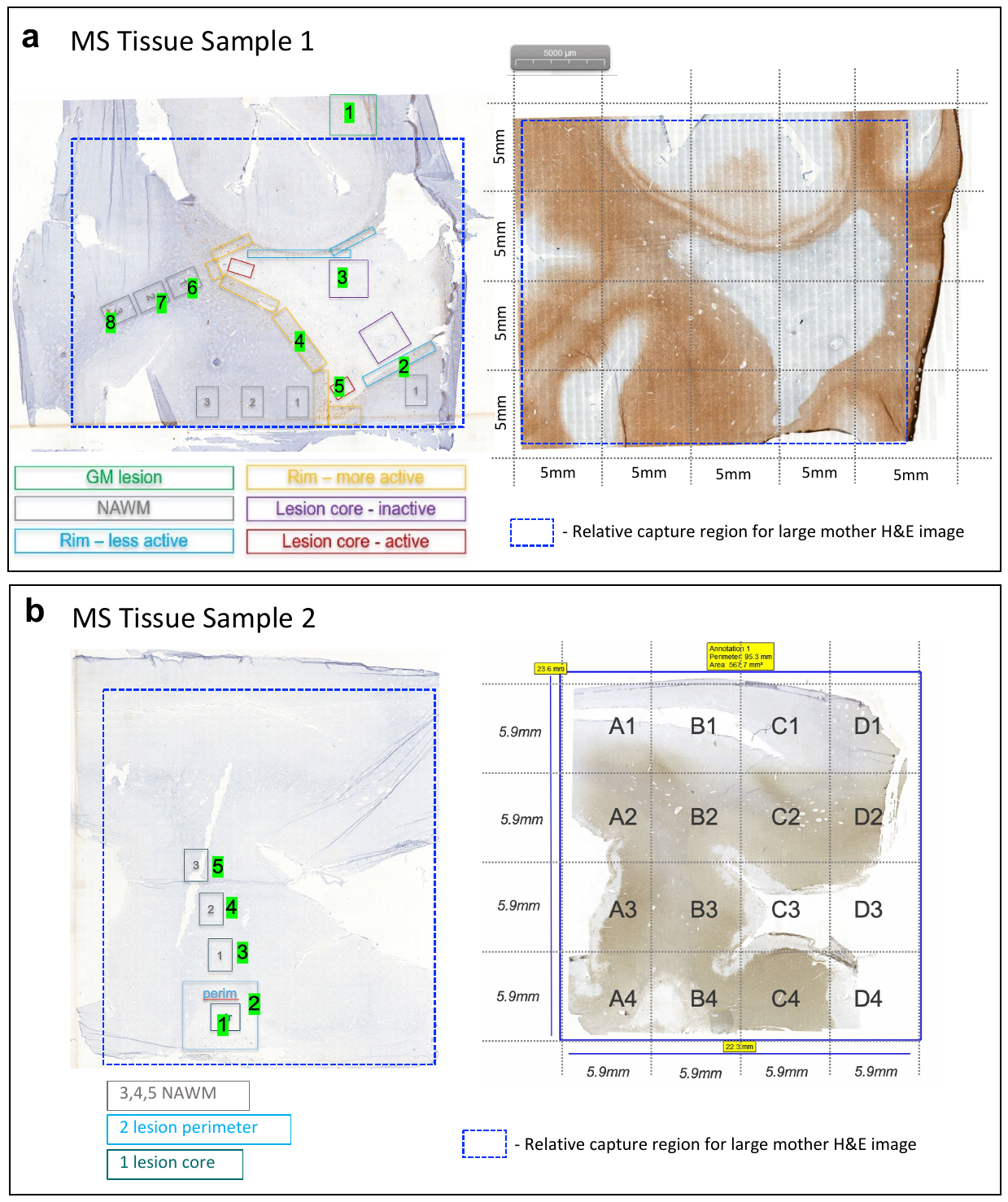

**Supplementary Fig. 10:** iSCALE removed gene expression batch effects in the multiple sclerosis Visium data. Left: before batch effect removal; Right: after batch effect removal.

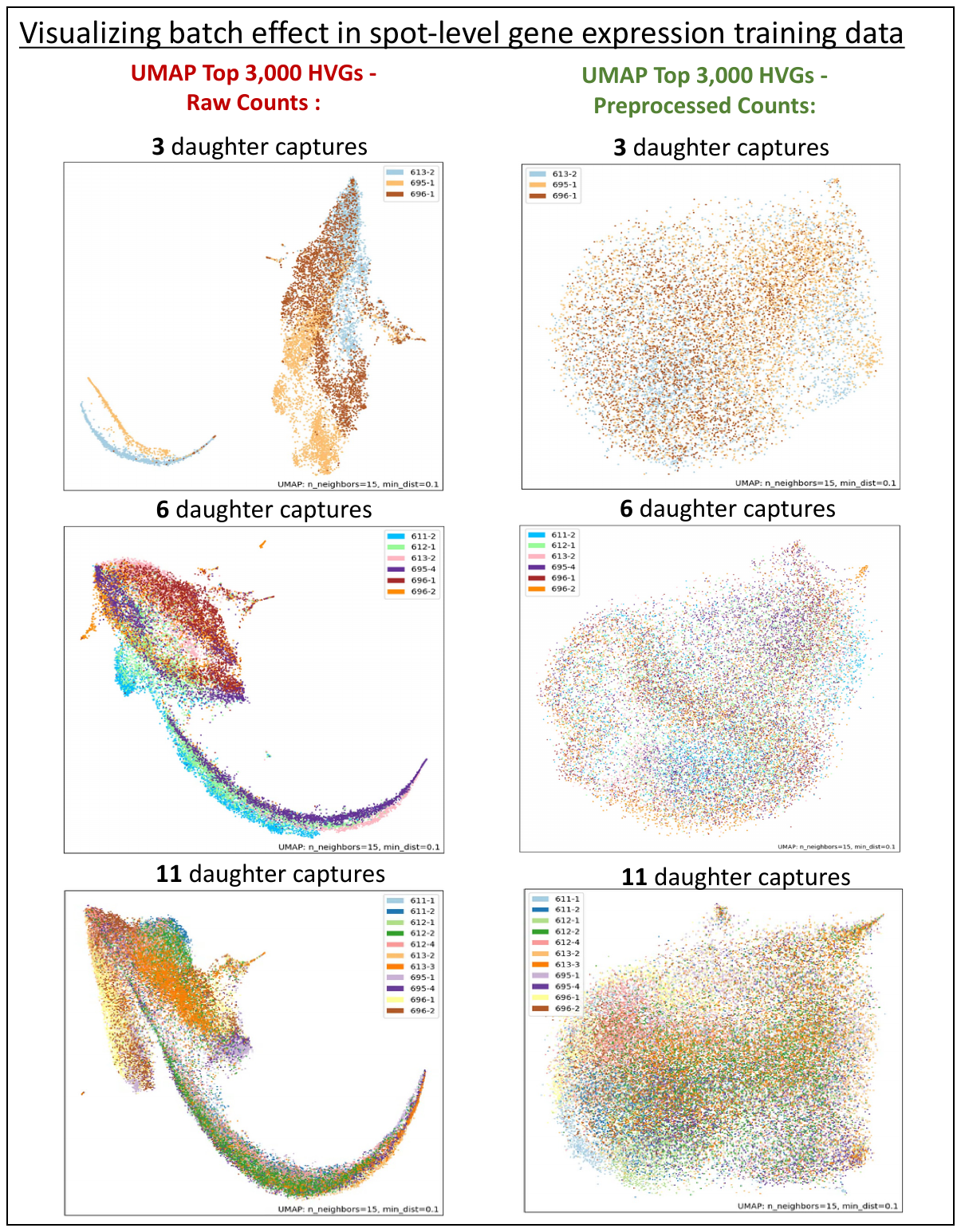

**Supplementary Fig. 11:** Visualization of iSCALE predicted gene expression in Sample 1 of the multiple sclerosis study.

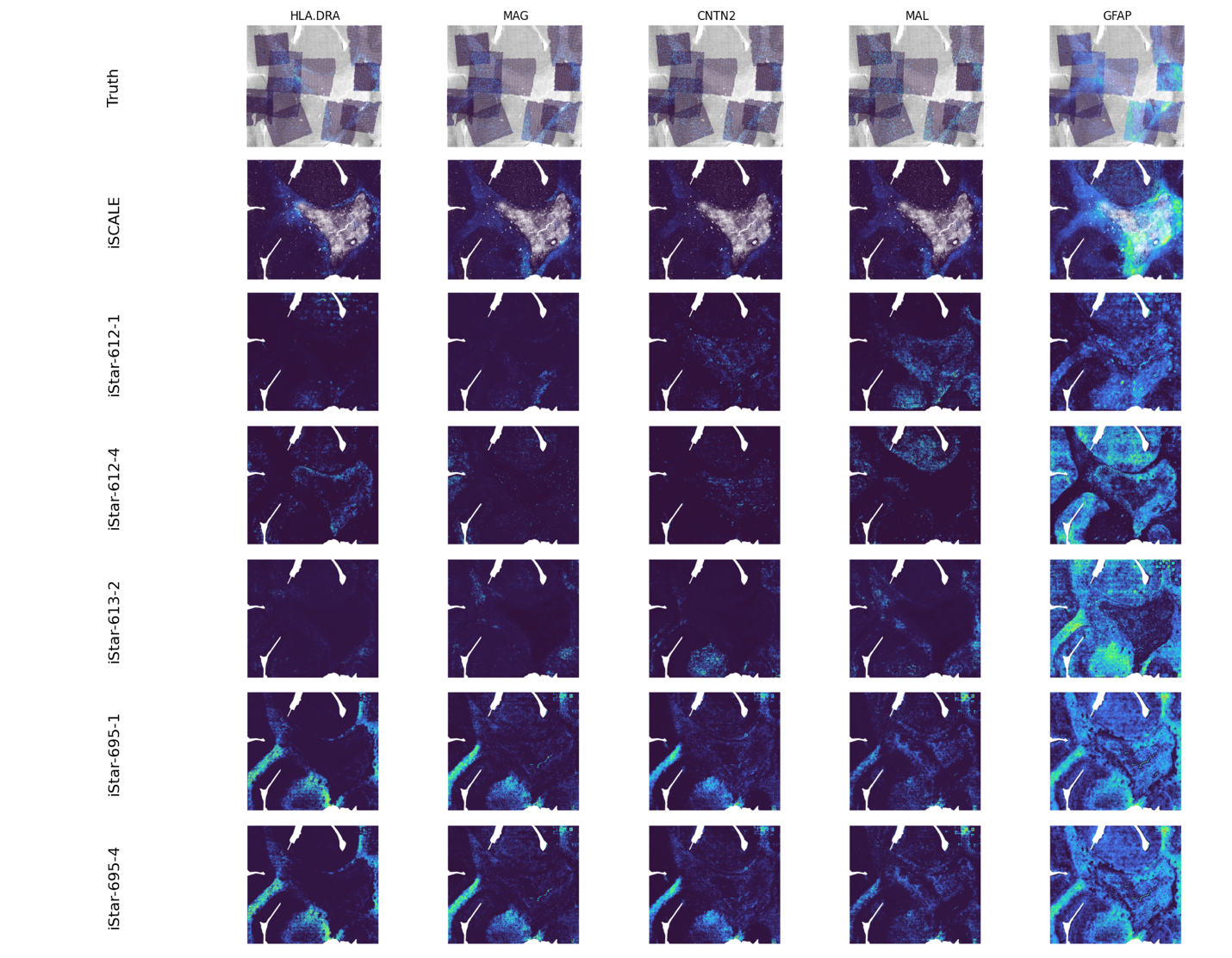

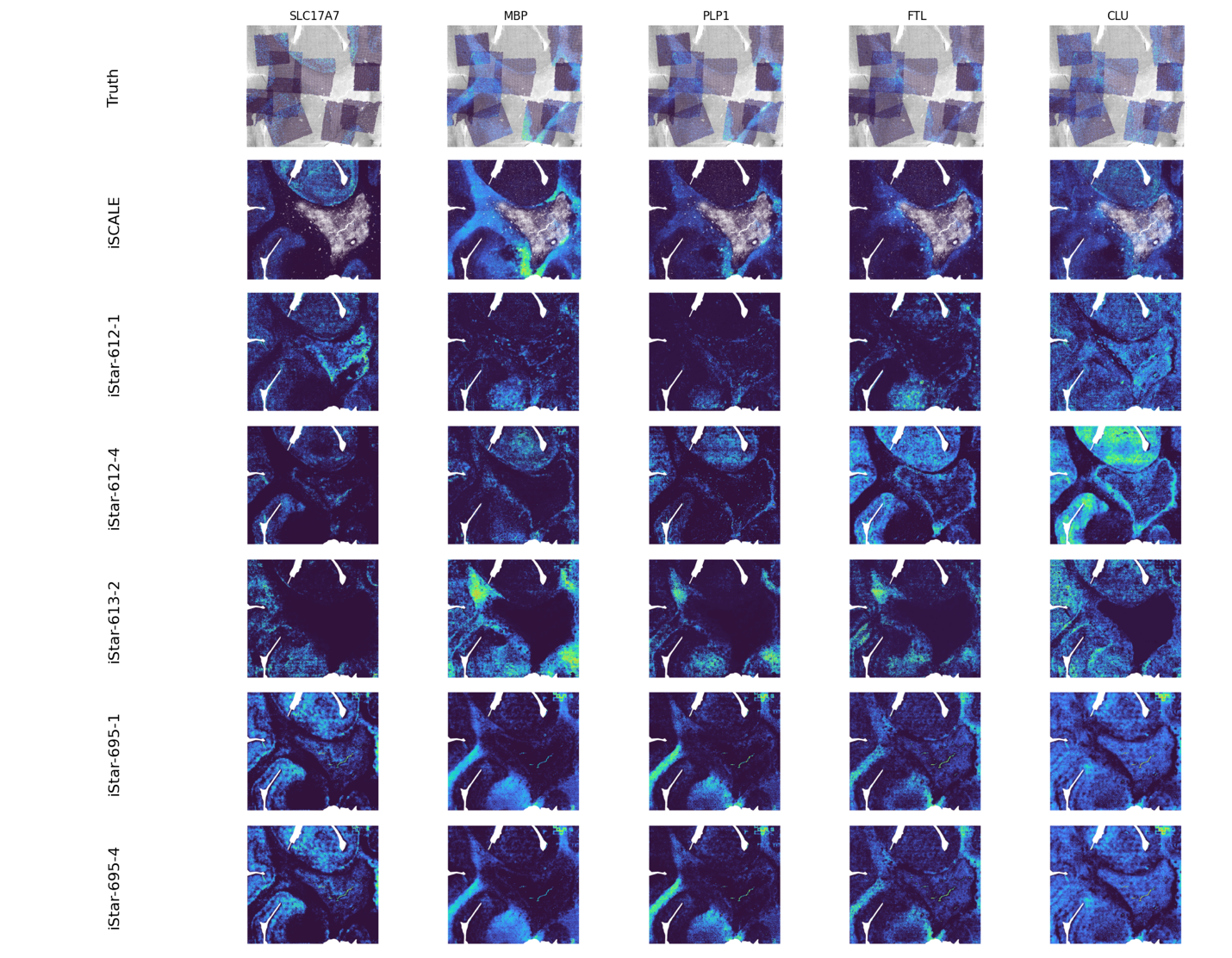

**Supplementary Fig. 12:** MS Sample 1 individual cell type masks based on iSCALE’s cell type annotation.

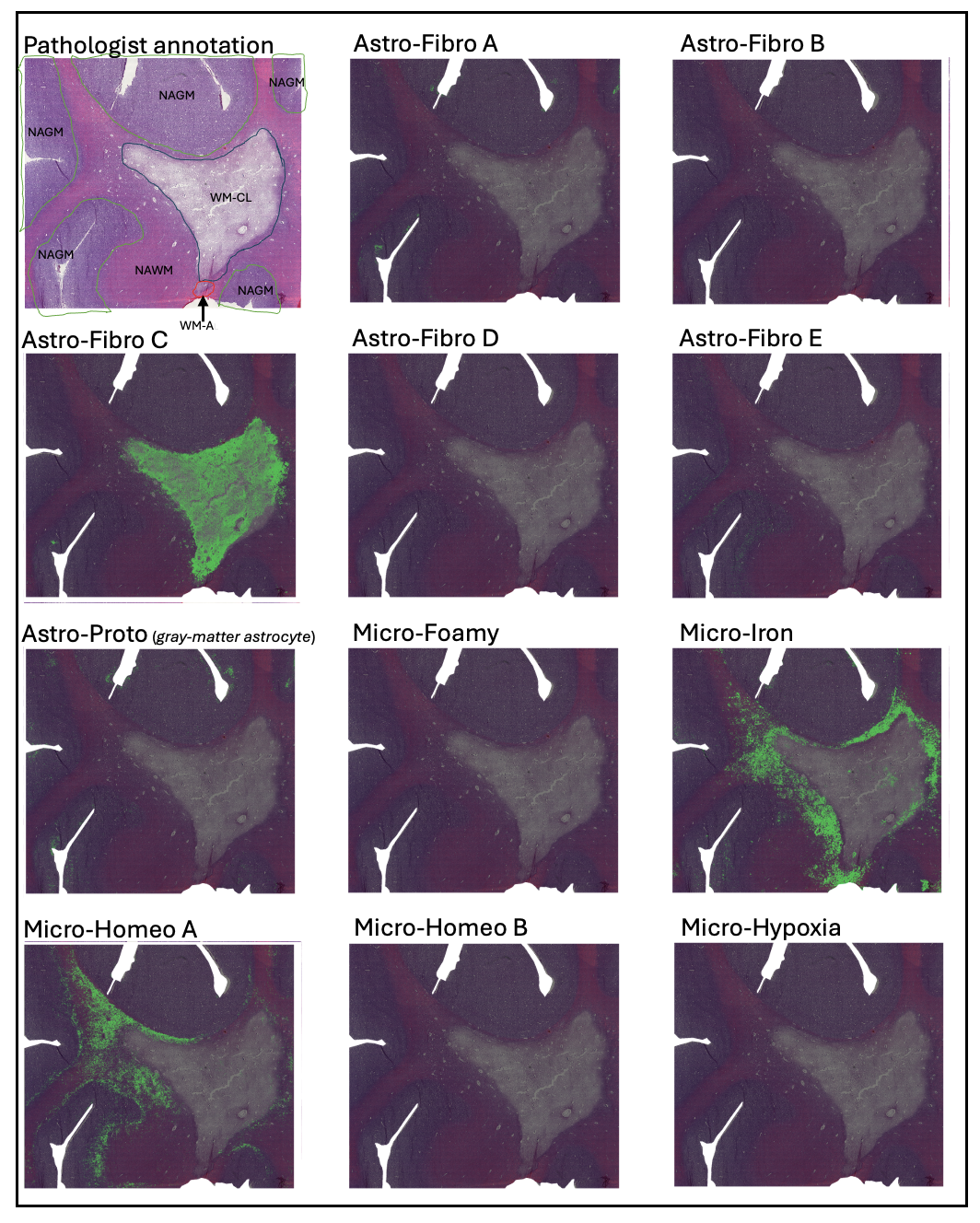

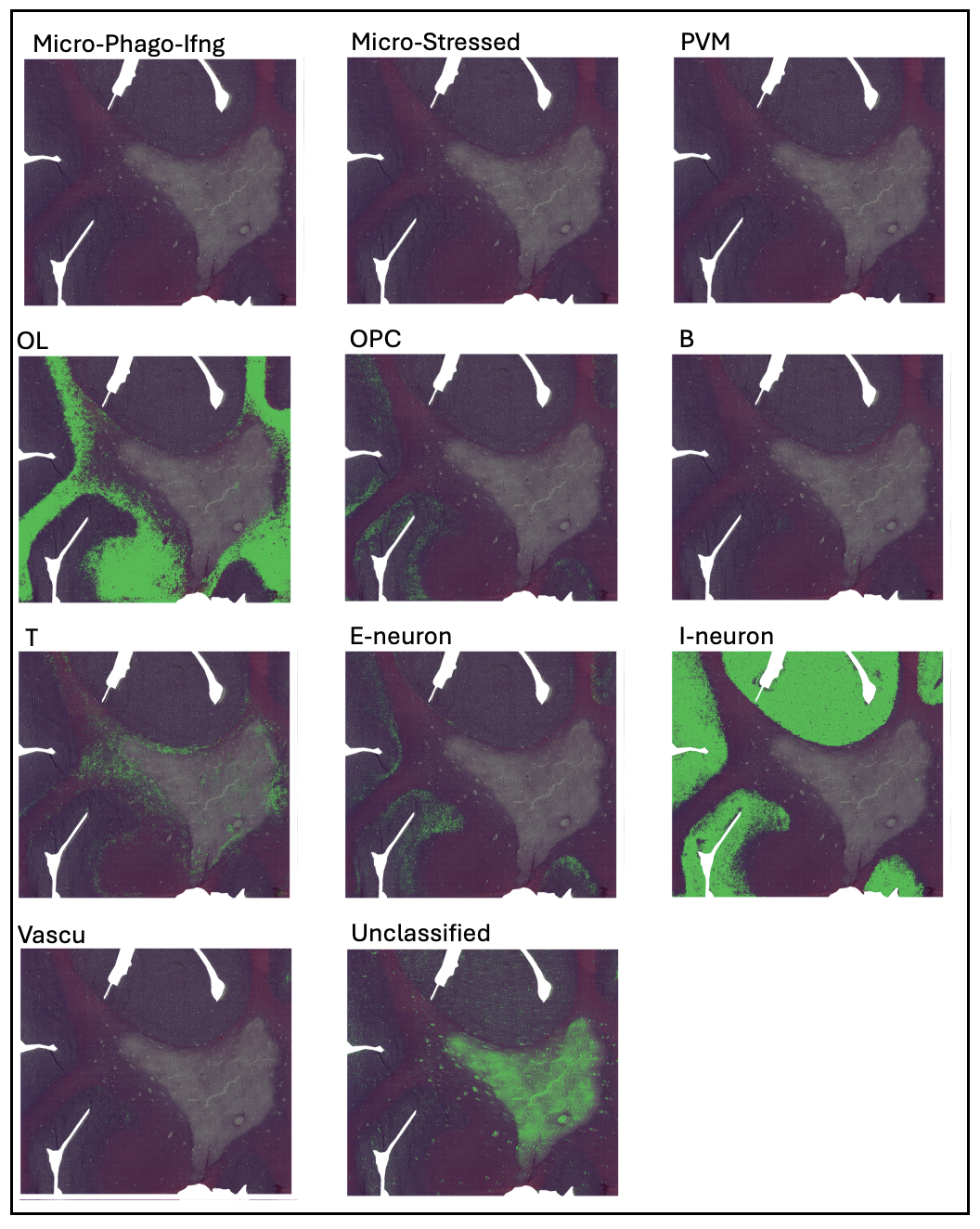

**Supplementary Fig. 13:** Cell type annotation by iStar for Sample 1 in the multiple sclerosis study.

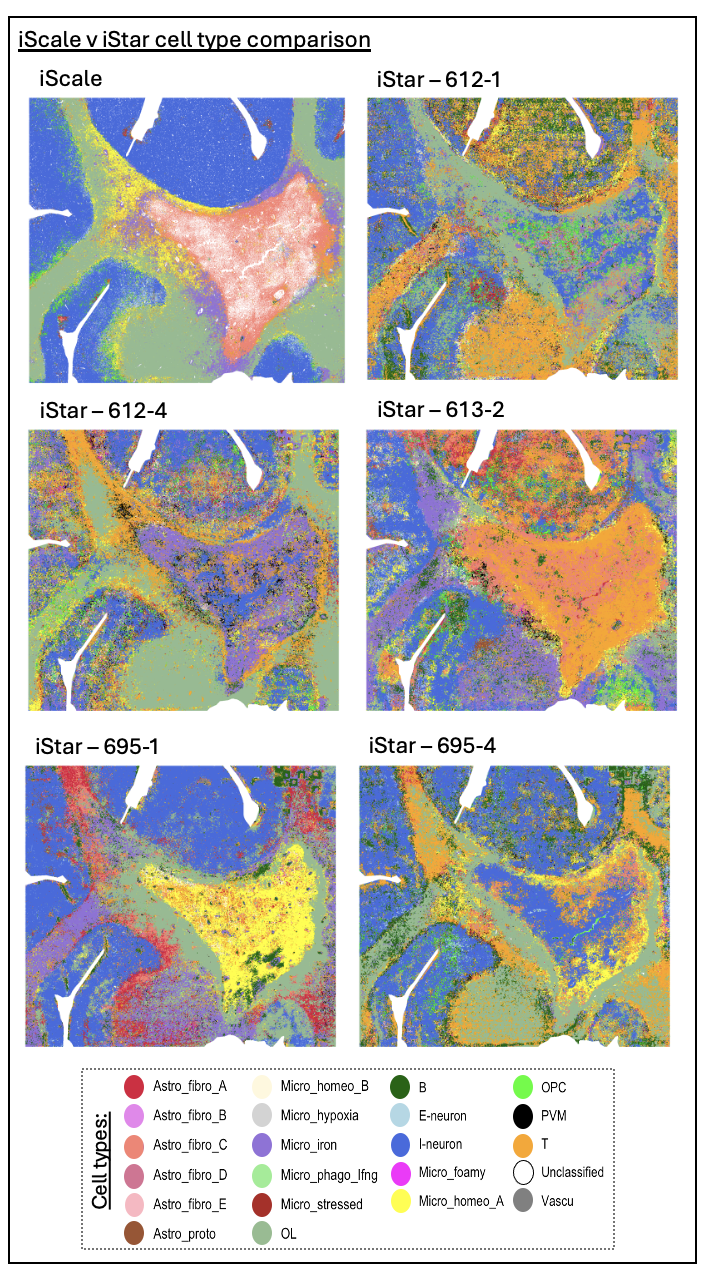

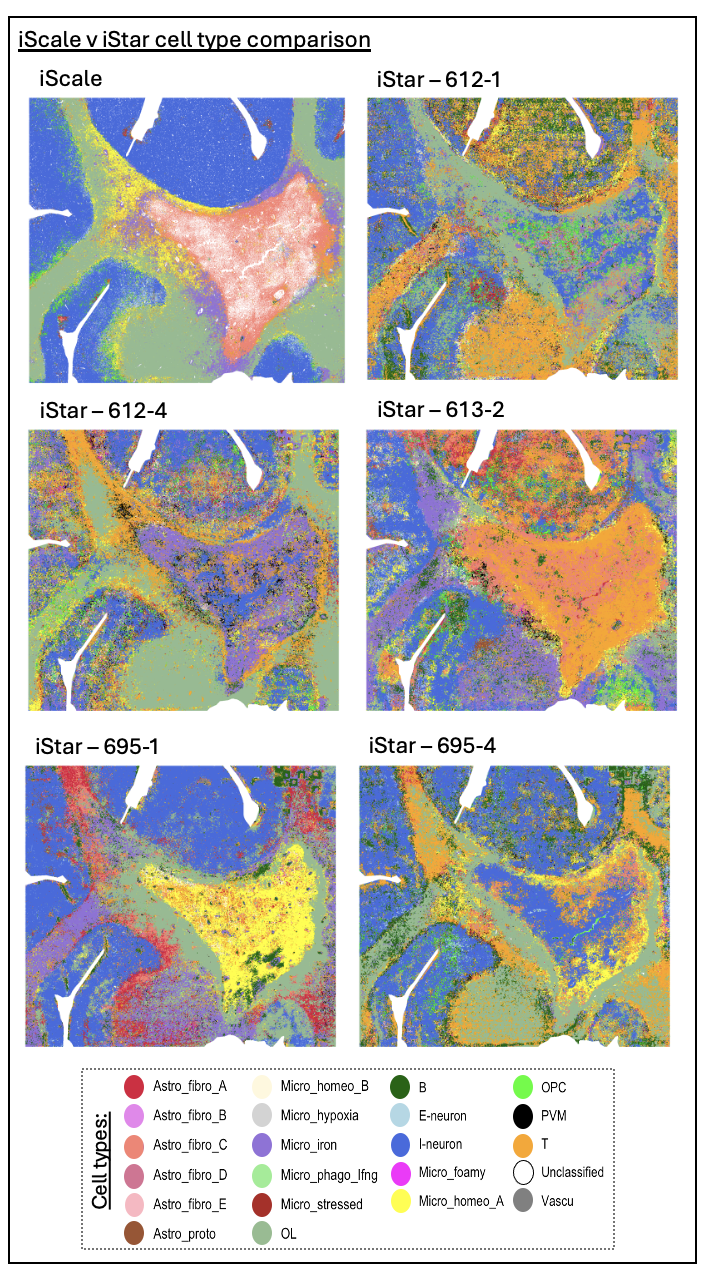

**Supplementary Fig. 14:** Out-of-sample cell type annotation results for individual cell-type masks for Sample 2 in the multiple sclerosis study.

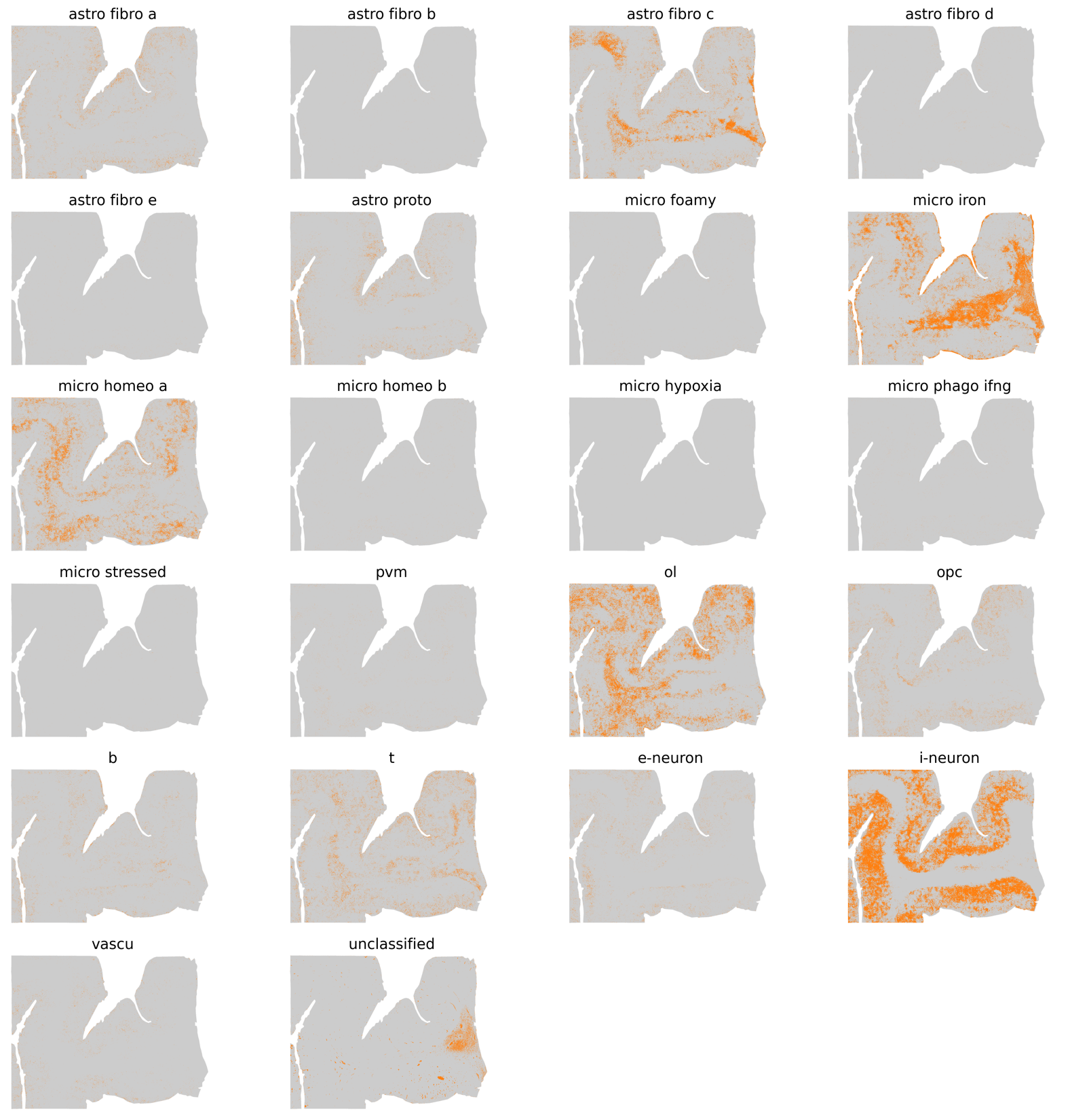

**Supplementary Fig. 15:** iSCALE segmentation of Sample 2 with varying cluster numbers in the multiple sclerosis study. Clustering results are shown for 4, 5, and 7 main clusters (cluster >1% of superpixels).

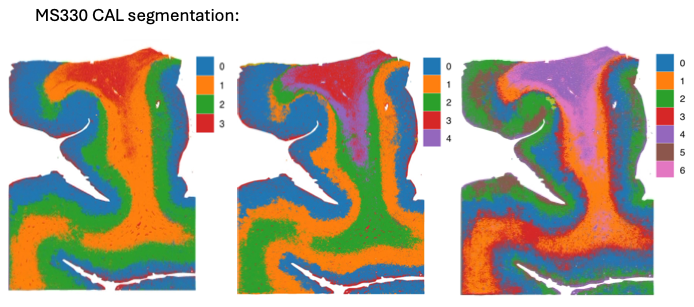

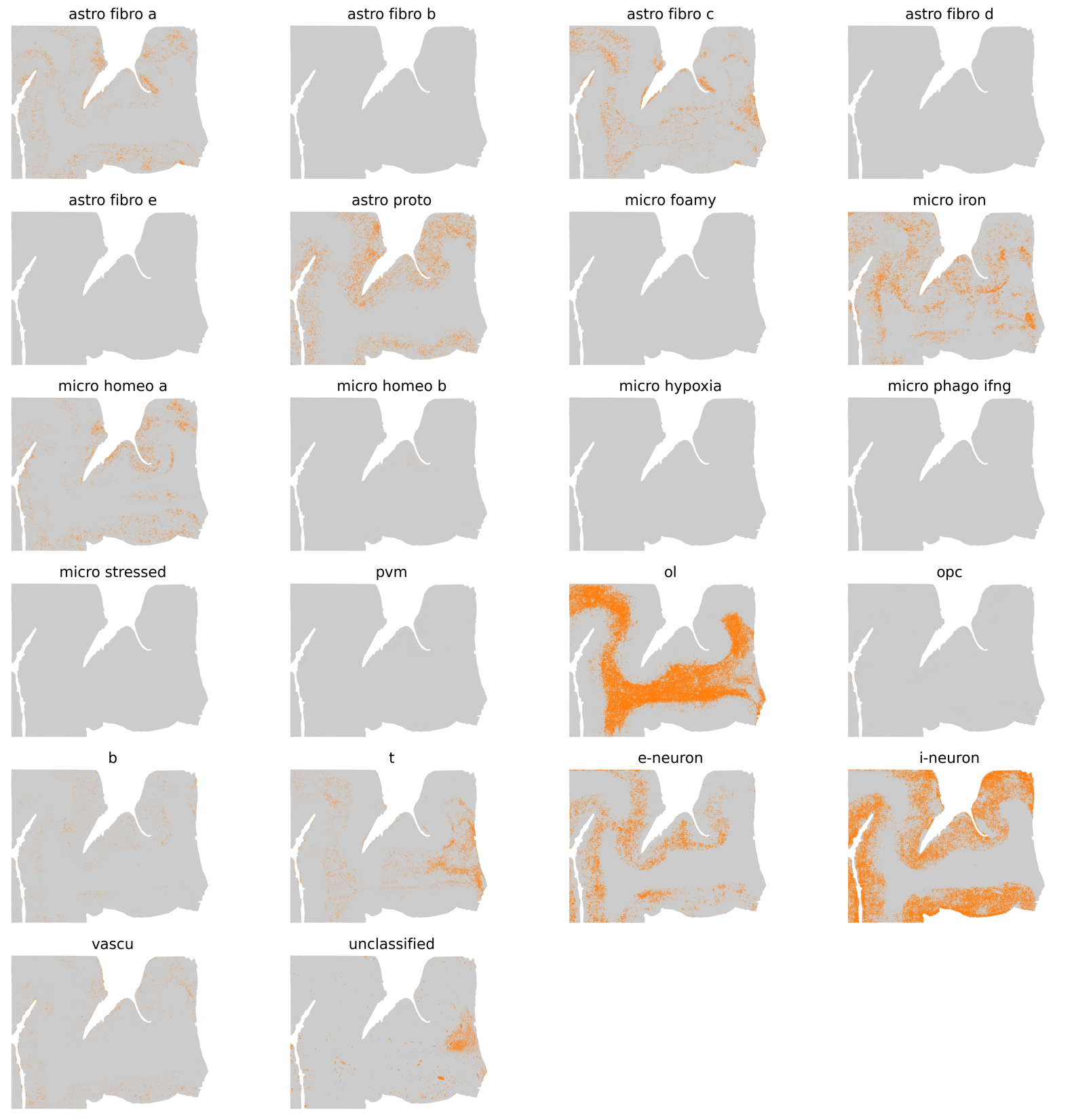
**Supplementary Fig. 16:** In-sample cell type annotation results for individual cell-type masks for Sample 2 in the multiple sclerosis study.

**Supplementary Fig. 17:** Detailed pathologist annotation of an adjacent tissue section to Sample 2 in the multiple sclerosis study.

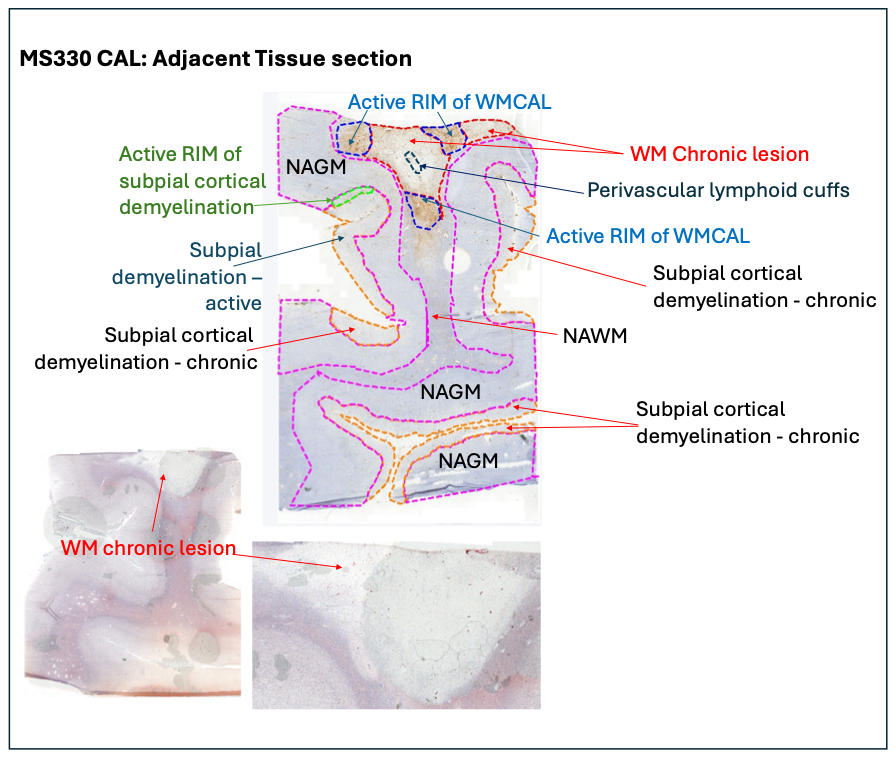

**Supplementary** **Table** **1**: Overview of Visium daughter captures generated in the MS study.

| Daughter Capture | Sample | Total number of genes | Number of non-zero genes | Number of spots |
| --- | --- | --- | --- | --- |
| 611-1 | sample 1 | 36,601 | 18,682 | 2,839 |
| 611-2 | sample 1 | 36,601 | 19,310 | 3,359 |
| 612-1 | sample 1 | 36,601 | 20,327 | 3,643 |
| 612-2 | sample 1 | 36,601 | 20,430 | 4,129 |
| 612-4 | sample 1 | 36,601 | 20,059 | 3,731 |
| 613-2 | sample 1 | 36,601 | 19,861 | 3,496 |
| 613-3 | sample 1 | 36,601 | 20,323 | 3,774 |
| 695-1 | sample 1 | 36,601 | 21,582 | 3,722 |
| 695-4 | sample 1 | 36,601 | 20,509 | 3,910 |
| 696-1 | sample 1 | 36,601 | 22,642 | 3,144 |
| 696-2 | sample 1 | 36,601 | 20,595 | 3,714 |
| A2 | sample 2 | 36,601 | 23,488 | 3,848 |
| A3 | sample 2 | 36,601 | 23,621 | 4,460 |
| A4 | sample 2 | 36,601 | 24,095 | 3,746 |
| B2 | sample 2 | 36,601 | 24,461 | 4,737 |
| B3 | sample 2 | 36,601 | 24,625 | 4,564 |
| B4 | sample 2 | 36,601 | 22,575 | 2,096 |
| **17 daughter captures** |  |  |  | **62,912 spatial observations** |

**Supplementary** **Table** **2**: Notation corresponding to the methods section of the manuscript, detailing the dimensions and providing a description for each referenced element.

| Notation | Dimension | Description |
| --- | --- | --- |
| $\boldsymbol{S}$ | *scalar* | total number of ST daughter captures used for training the iSCALE model |
| $\boldsymbol{q}_{\boldsymbol{s}}\boldsymbol{, s \in\{1,\ldots,S\}}$ | *scalar* | number of observed spatial observations (spots or cells) in daughter capture $s$ |
| $\boldsymbol{Q}$ | *scalar* | total number of observed observations across all $S$ daughter captures |
| $\boldsymbol{\phi}_{\boldsymbol{s}}\boldsymbol{, s \in\{1,\ldots,S\}}$ | $q_{s} \times2$ | matrix of 2D locations $(x, y)$of spatial coordinates for daughter capture $s$ |
| $\boldsymbol{L}$ | $Q \times2$ | combined matrix of spatial location across $S$ daughter captures |
| $\boldsymbol{G}_{\boldsymbol{s}}\boldsymbol{, s \in\{1,\ldots,S\}}$ | $q_{s} \times w_{s}$ | gene expression matrix for daughter capture $s$ |
| $\boldsymbol{w}_{\boldsymbol{s}}\boldsymbol{, s \in\{1,\ldots,S\}}$ | *scalar* | total number of genes measured for daughter capture $s$ |
| $\boldsymbol{\tau}_{\boldsymbol{s}}\boldsymbol{, s \in\{1,\ldots,S\}}$ | $\left\vert\tau_{s} \right\vert=w_{s}$ | set of all genes measured in daughter capture $s$ |
| $\boldsymbol{\varphi}$ | *scalar* | number of common genes among all S daughter captures |
| $\boldsymbol{\gamma}$ | $\left\vert\gamma\right\vert=\varphi$ | set of common genes among all $S$ daughter captures |
| $\boldsymbol{G}_{\boldsymbol{s}}\boldsymbol{, s \in\{1,\ldots,S\}}$ | $q_{s} \times\varphi$ | gene expression matrix for daughter capture$s$ subset to common genes |
| $\boldsymbol{G}$ | $Q \times\varphi$ | combined matrix containing genes expressions measures for all S captures for $\varphi$ common genes |
| $\mathfrak{G}$ | $Q^{*} \times\varphi^{*}$ | Smoothed gene expression matrix $G$ across adjacent daughter captures |
| $\boldsymbol{a}$ | *scalar* | number of highly variable genes to select, input parameter by the user |
| $\boldsymbol{K}$ | *scalar* | total number of genes used for model training |
| $\boldsymbol{℧}$ | $\left\vert℧ \right\vert=K$ | set of genes used in training the iSCALE model |
| $\mathfrak{G}^{\boldsymbol{*}}$ | $Q \times K$ | subset of matrix $\mathfrak{G}$to only the $K$ genes used for model training |
| ${\tilde{\mathfrak{G}}}^{\boldsymbol{*}}$ | $Q \times K$ | batch corrected normalized $\mathfrak{G}^{*}$ (measures for all $S$ captures for $K$ genes) |
| $\boldsymbol{X}$ | $X\in\mathbb{R}^{M}\times\mathbb{R}^{N}\times\mathbb{R}^{3}$ with height $M$ and width $N$ | the RGB-channel high-resolution mother histology image |
| $\boldsymbol{C}_{\boldsymbol{1}}$ | *scalar* | length of low-level local feature vector |
| $\boldsymbol{C}_{\boldsymbol{2}}$ | *scalar* | length of high-level global feature vector |
| $\boldsymbol{T}$ | $\left( M/256 \right)\times\left( N/256 \right)$ with $C_{1}\mathrm{channels}$ | high-level local feature image |
| $\boldsymbol{Z}$ | $\left( M/16 \right)\times\left( N/16 \right)$ with $C_{2}$ channels | low-level local feature image |
| $\boldsymbol{H}$ | $H\in\mathbb{R}^{M^{'}}\times\mathbb{R}^{N^{'}}\times\mathbb{R}^{C_{1}+C_{2}+3}$ with size $M^{'}\times N^{'}$ | combined histology feature image |
| $\boldsymbol{V}$ | *scalar* | total number of candidate cell types in gene marker set |
| $\mathcal{L}$ | *function* | weakly supervised loss function |
| $\boldsymbol{j}_{\boldsymbol{v}}\boldsymbol{, \upsilon\in\{1,\ldots,V\}}$ | *scalar* | number of marker genes for cell type $v$ |
| $\boldsymbol{\psi}_{\boldsymbol{v}}\boldsymbol{,}\boldsymbol{v}\boldsymbol{\in\{1,\ldots,V\}}$ | $\left\vert\psi_{v} \right\vert=j_{v}$ | list of marker genes names for cell type $v$ |
| ${\hat{\boldsymbol{Y}}}_{\boldsymbol{k}}\boldsymbol{, k \in\{1,\ldots,K\}}$ | ${(M}^{'}xN^{'})\times1$ | vector of predicted super-resolution gene expression |
| ${\tilde{\boldsymbol{Y}}}_{\boldsymbol{k}}\boldsymbol{, k \in\{1,\ldots,K\}}$ | ${(M}^{'}xN^{'})\times1$ | vector of predicted super-resolution gene expression standardized into the range of $[0.0, 1.0]$ using min-max approach |
| ${\tilde{\boldsymbol{Y}}}_{\boldsymbol{k}}^{\boldsymbol{*}}\boldsymbol{, k \in\{1,\ldots,K\}}$ | ${(M}^{'}xN^{'})\times1$ | vector of predicted super-resolution gene expression for gene $k$ standardized using percentile approach |
| $\boldsymbol{\theta}_{\boldsymbol{k}}$ | $K x 1$ | Vector containing training RMSE for each gene in the model |
| $\boldsymbol{P}_{\boldsymbol{95}}$ | *scalar* | Ninety-fifth percentile of training RMSE |
| ${\tilde{\boldsymbol{\Lambda}}}_{\boldsymbol{k}}$ | *scalar* | median super-resolution gene expression of gene k in training daughter captures |
| ${\tilde{\boldsymbol{\Phi}}}_{\boldsymbol{k}}$ | *scalar* | median super-resolution gene expression of gene k in mother capture |
| $\boldsymbol{F}_{\boldsymbol{k}}$ | $K x 1$ | Vector containing fold change between $\tilde{\Lambda}_{k}$and $\tilde{\Phi}_{k}$ for each gene in the model |
| $\boldsymbol{P}_{\mathbf{95}}^{\mathbf{*}}$ | *scalar* | Ninety-fifth percentile of $log(F_{k})$ |
| $\boldsymbol{\psi}_{\boldsymbol{v}}^{\boldsymbol{*}}\boldsymbol{,}\boldsymbol{v}\boldsymbol{\in\{1,\ldots,V\}}$ | $\left\vert\psi_{v}^{*} \right\vert=j_{t}^{*}$ | Refined list of marker genes names for cell type $t$ after the filtering method has been applied |
| $\boldsymbol{u}_{\boldsymbol{tmn}}$ | *scalar* | average marker score for superpixel $m,n$ for cell-type $t$ |
| $\boldsymbol{u}_{\boldsymbol{threshold}}$ | *scalar* | threshold used for assigning a cell-type label to a given superpixel |
